# Learning inhibitor interactions from protein deep mutational scanning data and deep learning models

**DOI:** 10.64898/2026.11.01.763444

**Authors:** Ziyue Zou, Isaiah Hazelwood, Gabriella Estevam, Jingyou Rao, Karson Chrispens, Matthew K. Howard, Willow Coyote-Maestas, James S Fraser

## Abstract

Recent advances in protein language models, geometric deep learning, and structure prediction frameworks have the potential to accelerate computational drug discovery. However, it remains unclear how well these methods generalize beyond their training sets. Mutation-dependent drug responses present a challenge to these methods because there are often large effects from relatively small perturbations. Here, we systematically benchmark diverse mutational effect and drug-target affinity prediction models using deep mutational scanning datasets spanning 11 inhibitor conditions for the MET kinase domain. Across both mutational effect prediction and drug–target affinity tasks, current AI models produce largely ligand-invariant predictions despite experimentally distinct inhibitor-dependent fitness landscapes. Ligand-aware geometric models, including LigandMPNN and LASErMPNN, partially recover experimentally validated resistance-associated hotspots, highlighting the importance of explicitly modeling protein–ligand interactions and local structural flexibility. In contrast, deep learning affinity prediction models underestimate mutation-induced changes in binding affinity. This deficiency likely reflects mutation-invariant structural priors derived from co-folded structure prediction frameworks and the absence of mutation-conditioned supervision in current affinity datasets. Finally, we further extended our analysis to several kinase-inhibitor deep mutational scanning datasets and confirm the overall trends. Our findings suggest that a primary limitation in improving performance on this task lies in the scarcity of mutation-conditioned protein-ligand training data rather than model architecture alone.

## Introduction

Computational approaches can substantially reduce the experimental burden of early-stage drug discovery by enabling in silico prediction of protein–ligand binding affinities. (Viet-Khoa Tran-Nguyen 2025; Walters and Murcko 2020; Schneider 2018; Vamathevan et al. 2019) Computer-aided drug design (CADD) has contributed to the development of therapeutics by providing atomistic insights into protein-ligand interactions, which can help rationalize structure-activity relationships (SAR) and guide the design of improved compounds (W. Yu and MacKerell 2017). A range of physics-based computational methodologies are widely employed to characterize protein-ligand interactions. These methods span molecular dynamics (MD) simulations, rigorous free energy methods such as free energy perturbation (FEP) (Zwanzig 1954; L. Wang et al. 2012) and thermodynamic integration (TI) (Kirkwood 1935; den Otter and Briels 1998), enhanced sampling techniques such as metadynamics (Laio and Parrinello 2002) and umbrella sampling (Torrie and Valleau 1977), as well as approximate endpoint approaches (e.g., MM-PBSA/GBSA) (Genheden and Ryde 2015). In addition, molecular docking methods (Shoichet et al. 2002) are commonly used for rapid screening and pose prediction, although their scoring functions generally provide only approximate estimates of binding affinity.

However, physics-based methods face several inherent limitations. In particular, MD simulations require sequential integration of Newton’s equations of motion, making them computationally expensive and difficult to scale. Their predictive accuracy also depends on factors such as force-field quality and accurate treatment of entropic contributions, both of which remain challenging. Advances in machine learning and computing hardware have enabled data-driven approaches to emerge as powerful alternatives. Classical machine learning models, including neural networks (Jain et al. 1996), random forests (Breiman 2001; Ballester and Mitchell 2010), and gradient-boosted trees (Chen and Guestrin 2016; J. Lu et al. 2019) have been widely applied to predict binding affinities and related biochemical properties from knowledge-based hand-crafted features defined *a priori*. (Rodrigues et al. 2021; Sun et al. 2021; Singh et al. 2025; Ballester and Mitchell 2010; Zilian and Sotriffer 2013; Svetnik et al. 2003; Durrant and McCammon 2011) More recently, deep learning (DL) methods have significantly advanced the related field by enabling high-throughput prediction and design across structural biology. These include protein structure prediction (Jumper et al. 2021)), mutational effect prediction using protein language models (C. Wang and Lu 2023; Lin et al. 2023; Bhatnagar et al. 2025; Frazer et al. 2021; Hesslow et al. 2022), drug-target affinity predictions (Jiang et al. 2026; R. Zhang et al. 2025; López-Ríos de Castro et al. 2026), and generative protein (Dieck-haus et al. 2024; Justas Dauparas et al. 2025; Fry et al. 2025; Birnbaum and Keating 2026; Ingraham et al. 2019; J. Dauparas et al. 2022; Ding et al. 2022; Scutteri et al. 2025; Kortemme 2024; Nijkamp et al. 2023) and small-molecule (Mistryukova et al. 2025; Özçelik et al. 2025; Atz et al. 2024; Xing et al. 2026; Bennett et al. 2023; Cremer et al. 2025; Dunn and Koes 2026; Sumita et al. 2026) design. In particular, AlphaFold (Jumper et al. 2021) and related structure prediction frameworks (Discovery et al. 2024; Passaro et al. 2025; Y. Zhang et al. 2026; Butcher et al. 2025; Krishna et al. 2024; L. Liu et al. 2024; Candido et al. 2026) have been extended to predict protein complexes and, more recently, protein-ligand co-folded structures, providing valuable structural priors for downstream CADD applications. These advances are especially impactful, as many computational drug discovery pipelines critically depend on accurate starting structures for downstream prediction of protein–ligand interactions.

Despite this success, accurately predicting the effects of single amino acid substitutions remains challenging in part because these perturbations induce subtle local changes that are underrepresented during pretraining (Feldman and Skolnick 2026; Huynh et al. 2026). Protein language models (pLMs), particularly single-sequence pLMs, offer an attractive alternative for modeling mutational effects at scale. (Elnaggar et al. 2020; Brandes et al. 2022; Bepler and Berger 2019; Ferruz et al. 2023) Among these, the Evolutionary Scale Modeling (ESM) (Lin et al. 2023; Hayes et al. 2025; Rives et al. 2021; Candido et al. 2026) family has demonstrated strong performance in zero-shot prediction of residue-level mutational effects, typically using log-likelihood ratio between wild-type and mutant sequences. The resulting representations have also proven effective for supervised transfer learning across a wide range of downstream tasks. (Ouyang-Zhang et al. 2025; Dinh et al. 2026) Recent work has further extended protein representations to incorporate more expressive tokenizations, including structure-aware and multi-scale representations. (Su et al. 2023; M. Li et al. 2024; Tan et al. 2025; Cheng et al. 2023; Ruffolo et al. 2024; Jantos et al. 2022). The mechanisms underlying the excellent performance of pLMs in mutational prediction remain an active area of investigation, with proposed explanations ranging from learned evolutionary information (Hayes et al. 2025), interacting sequence motifs (Z. Zhang et al. 2024), in-context positional effects (Vieira et al. 2026), or learned representations corresponding to structural, functional, and protein-family-level biological features (Parsan et al. 2025; Simon and Zou 2025; Gujral et al. 2025; Adams et al. 2025; Candido et al. 2026). In parallel, a growing class of transformer-based models has been developed for small molecules, encoding ligand information directly from Simplified Molecular Input Line System (SMILES) (Weininger 1988) strings or molecular graphs. (Chithrananda et al. 2020; Irwin et al. 2022; Ross et al. 2021; Born and Manica 2023) Together, these developments provide the foundation for integrated modeling of protein–ligand interactions using learned representations of both proteins and small molecules.

Many modern drug-target affinity models integrate representations learned by pretrained protein and ligand encoders. (Jiang et al. 2026; Vefghi et al. 2025) These models combine learned embeddings using fusion strategies, including simple concatenation (Öztürk et al. 2018), attention-based mechanisms such as bi-directional attention (Huang et al. 2022) and cross-attention (X. Zhang et al. 2024; Kawaharada et al. 2026), as well as higher-order interactions through bilinear attention (Bai et al. 2023). These approaches typically operate on sequence- or graph-level representations and aim to predict scalar binding affinities without explicitly modeling binding geometry. When protein-ligand structures are available, whether from experiments, simulations, or co-folding prediction, geometry-based approaches can provide a more physically grounded alternative. In this setting, interactions are modeled explicitly using spatial representations such as binding pocket graphs or equivariant neural networks that capture atomic-level interactions (Z. Xie et al. 2025; Graber et al. 2025; J. Liu et al. 2026; Sim and Lee 2026; J. Yu et al. 2026) More recently, diffusion and flow-matching models have emerged as powerful generative frameworks for protein-ligand docking. These methods iteratively refine ligand poses (and in some cases protein conformations) through learned stochastic or deterministic dynamics, ultimately generating bound complexes that can be used for downstream binding affinity estimation. This paradigm offers a flexible alternative to traditional docking approaches by directly modeling the distribution of plausible binding configurations. (Corso et al. 2022, 2024; Passaro et al. 2025; Cremer et al. 2025; Morehead and Cheng 2024; W. Lu et al. 2022; Stärk et al. 2022)

Although protein-ligand affinity datasets span diverse protein families and chemical scaffolds, it is unclear how well predictive methods generalize. Contamination in training abounds for deep learning methods and the physics-based methods are rarely tested prospectively at scale. One potential source of information that has not been leveraged is deep mutational scanning data in the presence of different chemicals. Deep mutational scanning (DMS) provides a complementary experimental framework for evaluating model generalization by systematically measuring the phenotypic consequences of large numbers of sequence variants in parallel (Fowler and Fields 2014). Unlike conventional affinity benchmarks that span many unrelated proteins and ligands, DMS densely samples sequence space around a common protein background, often at single-amino-acid resolution. When the same mutational library is screened under multiple inhibitor conditions, it additionally measures how the functional consequence of an identical mutation changes across chemical environments. These effects emerge from the interplay between sequence variation, protein structure, and protein-ligand interactions. Although measured indirectly through fitness assays, the differential fitness relative to an unperturbed control experiment can be used as a proxy for affinity. More importantly, systematic matched measurements of wild-type and single-mutant protein-ligand interactions are sparse in commonly used affinity datasets and may be further depleted by sequence-identity-based filtering or clustering procedures designed to prevent train–test leakage. Protein language models have demonstrated strong zero-shot performance in predicting mutational effects across diverse protein fitness assays. However, this performance primarily reflects prediction of mutation-dependent fitness in a fixed experimental context; whether the same representations generalize to changes in mutational effects across ligand environments remains unclear (Hayes et al. 2025; Z. Zhang et al. 2024; Vieira et al. 2026; Parsan et al. 2025; Simon and Zou 2025; Gujral et al. 2025; Adams et al. 2025).

To investigate mutation- and ligand-dependent functional effects, we evaluated several state-of-the-art (SOTA) deep learning models on a recent Deep Mutational Scan (DMS) (Fowler and Fields 2014; Araya and Fowler 2011) dataset measuring the effects of single-residue mutations within the kinase domain of hepatocyte growth factor receptor (MET), a clinically important drug targets (Ou et al. 2024; Jiao et al. 2018), across 11 inhibitor conditions. (Estevam et al. 2025; Rao et al. 2026, 2024) This dataset captures functional changes arising from the interaction between single-point mutations and inhibitor environment. We benchmarked models originally developed for either mutational effect prediction or drug-target affinity estimation to assess their ability to predict functional changes under combined mutation and ligand perturbations. Using activity splitting, we further analyzed model performance in identifying sensitizing and drug-resistance mutations across different inhibitor conditions. To evaluate the generalizability of these observations, we expanded the benchmark to additional kinase–inhibitor datasets involving FGFRs (Tangermann et al. 2025), EGFR (Kim et al. 2024), SRC (Chakraborty et al. 2024), and CDKs (Persky et al. 2020; Gould et al. 2025). Collectively, this framework enables a systematic assessment of how current deep learning models capture mutation-dependent protein–ligand interactions and provides guidance for the development of more robust predictive models for drug response.

## Results

### MET DMS data for mutant-specific ligand binding estimates

We first characterized the MET deep mutational scanning (DMS) dataset used for benchmarking deep learning models by examining the distribution of variant fitness scores across 11 inhibitor conditions (**Figure 1**A). Relative to the dimethyl sulfoxide (DMSO) control, individual inhibitors produced distinct shifts (computed by mean absolute bin migration, *⟨|*Δ*b|⟩*, relative to the DMSO control) in the mutational fitness landscape, indicating substantial ligand-dependent differences in the functional consequences of single-residue substitutions. Although all inhibitors bind within or adjacent to the adenosine triphosphate (ATP)-binding pocket of MET, they stabilize different kinase conformations (Estevam et al. 2025). Type I inhibitors bind the active DFG-in conformation and, more subtly, they can be classified into type Ia or Ib by whether the molecules interact with the solvent front residue G1163 (Jean Cui 2013). Type II inhibitors extend into the hydrophobic back pocket and stabilize an inactive kinase state by disrupting the regulatory R-spine. Type I½ inhibitors combine features of both binding modes, whereas type III inhibitors bind an allosteric site without directly competing with ATP. Notably, the type III inhibitor tivantinib produced the smallest overall shift in variant fitness relative to the DMSO control (**Figure 1**A), consistent with its limited effect on suppressing MET mutant enrichment. We therefore retained tivantinib in the benchmark because it provides a useful reference for evaluating whether models distinguish ligand-dependent functional effects across mechanistically distinct inhibitor classes.

**Figure 1.**
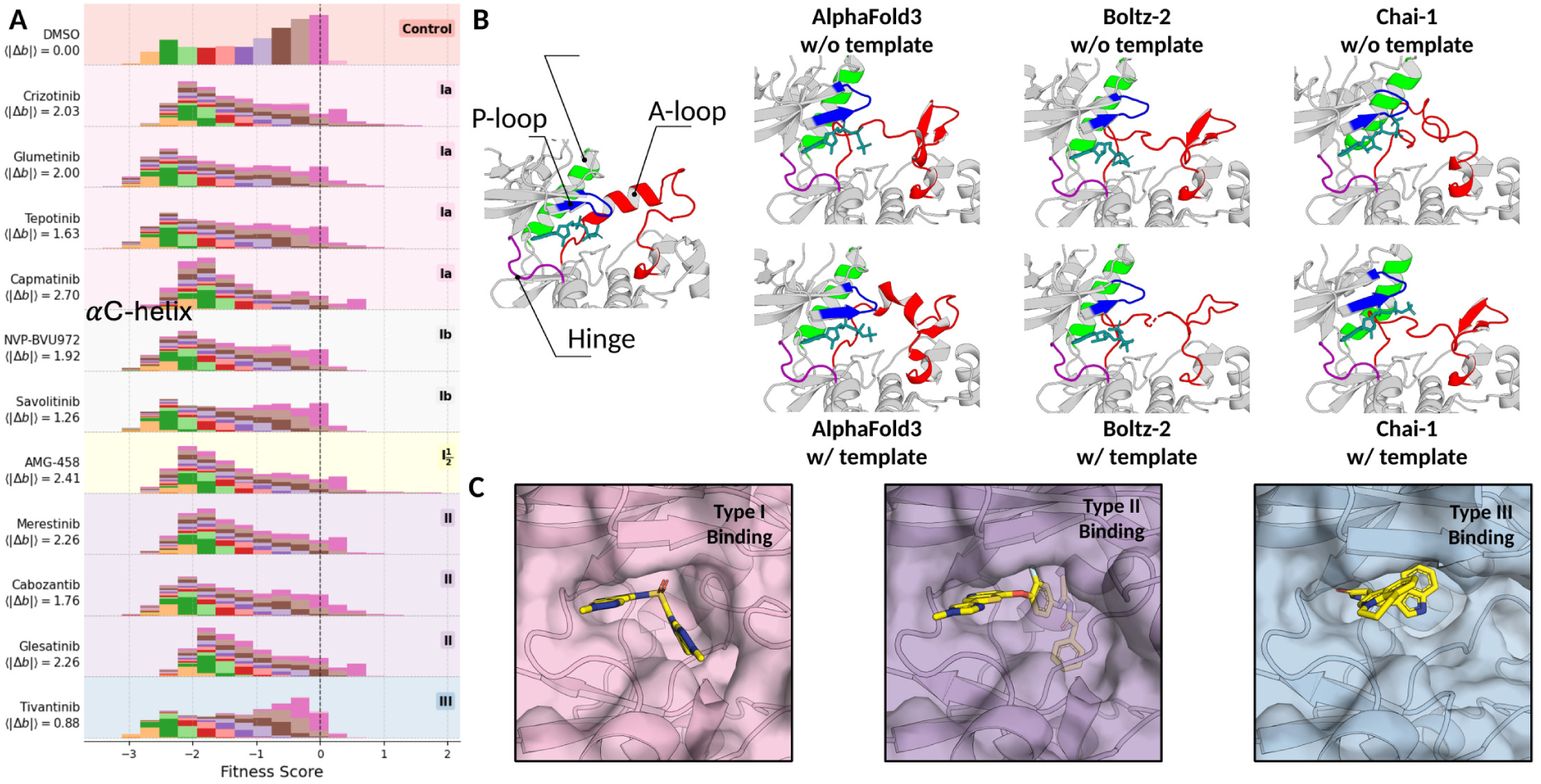
MET DMS fitness landscapes across inhibitors and predicted MET kinase complexes. A) Experimental measured fitness scores in treatments. Variants within the same bin in control condition are color coded and traced for other conditions. For each condition, the averaged movement of variants from bin-to-bin (Δb) is computed with respect to the DMSO control to show the effects of the inhibitors. The magnitude of these distributional shifts did not strictly follow inhibitor class, suggesting that chemically related inhibitors can induce distinct mutational landscapes. B) MET kinase-ATP complexes predicted by different deep learning models - AlphaFold3, Boltz-2 and Chai-1 with and without template matching. In particular, key functional regions are colored - P-loop (in blue), hinge (in purple), αC-helix (in green), and A-loop (in red). C) Cofolded structures of MET kinase upon binding to kinase inhibitors of different types predicted by AlphaFold3.

### Structure predictions on cofolding complexes

To compare across protein-ligand affinity prediction models, we needed to generate structures of the relevant protein-ligand complexes for those models reliant on structural inputs. Because many experimentally determined protein structures deposited in the Protein Data Bank (PDB; (Berman et al. 2000)) contain unresolved residues or partially resolved domains, we evaluated whether deep learning-based structure prediction could generate complete protein– ligand complexes suitable for this task. Specifically, we predicted ATP-bound structures of the MET kinase domain to represent its experimental conformation under DMSO control condition using three state-of-the-art co-folding models: AlphaFold3 (Abramson et al. 2024), Boltz-2 (Passaro et al. 2025), and Chai-1 (Chai Discovery et al. 2024). For each, we produced predictions with and without template information. All predicted complexes aligned well with the experimentally determined reference structure (PDBID 3DKC; (Buchanan et al. 2009), **Figure 1**B), with template guidance consistently improving reconstruction accuracy. AlphaFold3 achieved the highest structural accuracy, reducing the global backbone RMSD from 3.7 Å without templates to 1.2 Å with template matching. Boltz-2 and Chai-1 also benefited from template guidance, with backbone RMSDs improving from 2.8 to 2.5 Å and from 4.0 to 3.0 Å, respectively. The large deviation in backbone RMSDs is likely due to the conformation change in A-loop where the deposited structure introduced two stabilizing mutations and the structure predictions were made with the wildtype sequence (**Figure 1—figure supplement 1**). Although the predicted ligand poses differed among models along with the A-loop region, ATP remained deeply buried within the binding pocket in all cases, consistent with previous observations that current co-folding models memorize binding pocket locations and do not always recapitulate detailed interactions (Masters et al. 2025). These results indicate that AI-predicted co-folded structures have the potential to provide structural inputs for evaluating geometry-aware models in subsequent analyses for ligands, like ATP, that are in the training set, but that such models may not be as accurate as experimentally determined structures.

We next generated predicted MET kinase complexes for all 11 inhibitor conditions using the three structure prediction frameworks described above (**Figure 1**C). Because experimentally resolved template structures were unavailable for some inhibitor-bound complexes, all structures were predicted without template guidance and subsequently used as inputs for the structure-aware models evaluated in this study. The unavailable 3-dimension structures in PDB are cabozantinib (type II), glesatinib (II), capmatinib (Ib), and glumetinib (Ib). For the other complexes, we note that the specific ligand-bound structures were available in the training sets for these predictors. To validate whether structures predicted with or without template guidance will affect models’ predicted fitness landscape, we evaluated several complexes whose structures are available in PDB. Our results suggest the predicted fitness landscapes among these complexes did not differ significantly between structures predicted with template-free and template-guided (**Figure 1—figure supplement 2**) and indicating these structure-based models are robust to the structure generation protocol. We therefore used the template-free predicted complexes from identical inference settings for all subsequent analyses which provides a consistent basis for downstream benchmarking.

### Models without explicit ligand representations do not predict ligand-specific mutational fitness

Having established a consistent set of predicted structures across inhibitor conditions, we next asked whether sequence and protein structural information alone are sufficient to recover ligand-dependent mutational fitness. To establish this baseline, we evaluated three representative zero-shot mutational effect predictors: the sequence-only protein language model ESM, the structure-based geometric model ProteinMPNN, and the energy-based diffusion model ProteinEBM (Roney et al. 2025) (**Figure 2**A). ESM and ProteinMPNN models estimate mutational effects from changes in log masked marginal probabilities between wild-type and mutant residues, computed from either sequence representations (ESM) or protein graph representations (ProteinMPNN). ProteinEBM is a more recent approach that scores conformations on free-energy landscapes using diffusion-based generative modeling. Importantly, ESM and ProteinEBM receive identical protein sequences for every inhibitor condition and therefore cannot explicitly encode ligand-dependent effects. In contrast, for ProteinMPNN, we input AlphaFold3-predicted protein conformations generated for each inhibitor-bound complex. In this manner, ProteinMPNN does not explicitly account for interactions with the ligand for its message passing and consequential scoring steps. However, the ligand interactions are accounted for in the input model and can therefore indirectly change the probabilities for each amino acid. Under the DMSO control condition, the ESM2 model achieved moderate agreement with experimentally measured fitness as our baseline method (Spearman’s ρ ≈ 0.50). However, its performance decreased by an average of 53% across inhibitor-treated conditions, indicating limited sensitivity to ligand-dependent functional changes despite strong baseline performance. ProteinEBM showed comparable performance to the MPNN models under both untreated and inhibitor-treated conditions with a smaller average decline in predicting the inhibitor-treaded conditions with average decreases of 43%. This may suggest that ProteinEBM can identify stable variants that are insensitive to inhibitor conditions. Protein-MPNN also exhibited a large reduction in performance (a decrease of 54%) despite receiving distinct inhibitor-specific structural inputs, suggesting that differences in predicted protein conformations alone are insufficient to recover ligand-specific mutational effects. Collectively, these results suggest that although zero-shot mutational effect predictors do well on predicting protein fitness in the unperturbed condition, they cannot predict the effects of ligand perturbations because the main driving force, protein-ligand interactions, remains missing from their input.

**Figure 2.**
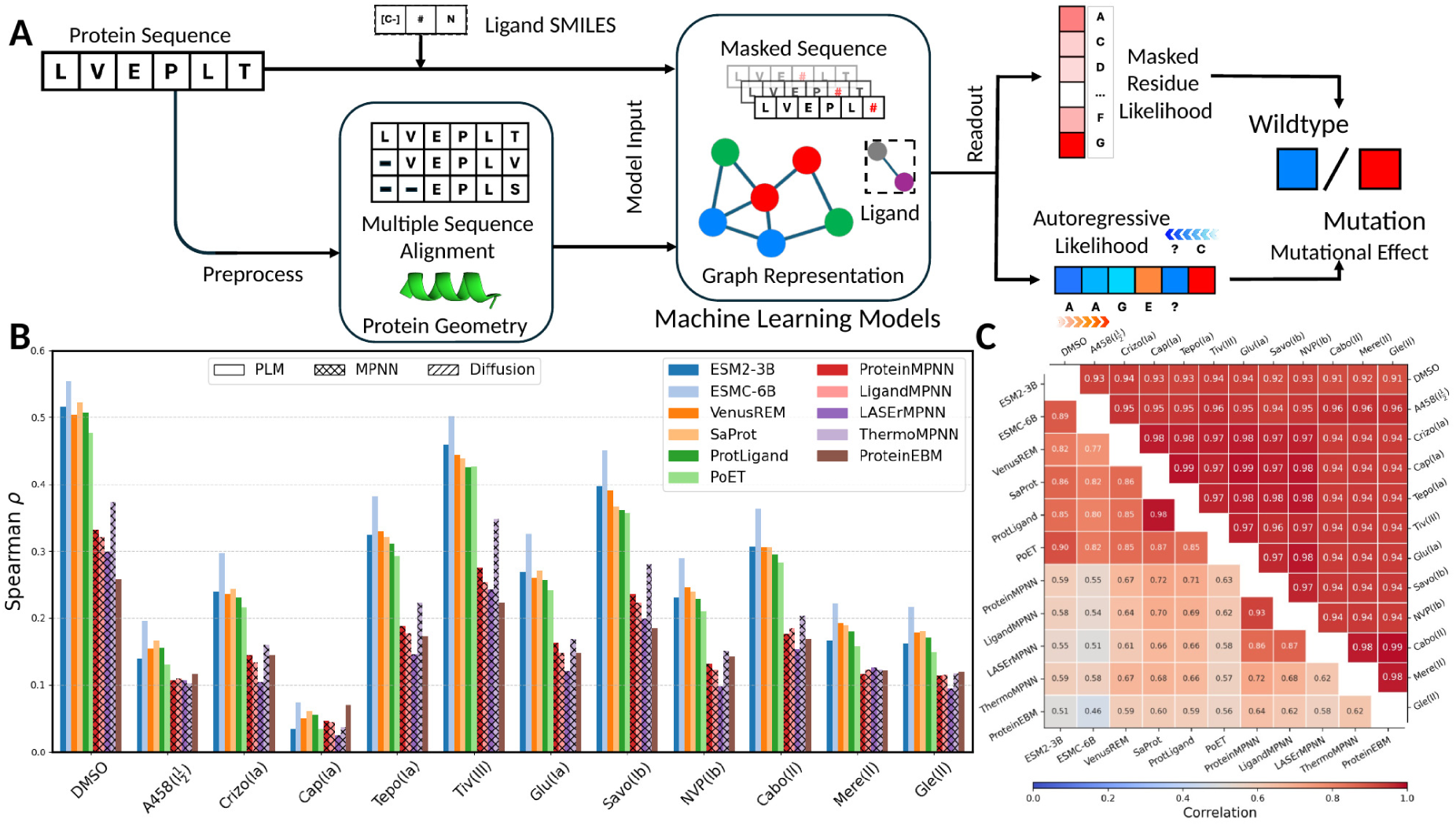
Mutational effect predictions from models without explicit ligand representations. A) A summary of workflow of current deep learning models on predicting protein mutational effects. B) Bar plot on Spearman correlation coefficients of model predictions to experimental measured fitness scores. C) Averaged Pearson correlation coefficients between mutational effect predictions across treatments of all models (upper triangle) and mean Spearman correlation coefficients between model mutational effect predictions over contexts (lower triangle).

We next evaluated recently developed protein language models that augment sequence representations with structural information (Su et al. 2023; M. Li et al. 2024; Ouyang-Zhang et al. 2025), evolutionary context from multiple sequence alignments (Jantos et al. 2022), or both (Tan et al. 2025). In addition, we assessed recent graph- and geometry-based generative models originally developed for protein design (Birnbaum and Keating 2026) and subsequently adapted for mutational thermostability prediction (Dieckhaus et al. 2024). Despite these architectural differences, mutational effect predictions improved by less than 0.05 in Spearman correlation compared to the baseline ESM2-650M model in the control condition (**Figure 2**B and **Figure 2—figure supplement 1**A) and the next generation ESM model (Candido et al. 2026), ESMC, outperforms all these PLM variants. Similar to the baseline, all models showed performance reductions of approximately 50% under inhibitor-treated conditions relative to the DMSO control. Notably, providing full-length protein sequences, and corresponding predicted full-length structures when required, did not consistently improve mutational effect prediction compared with using the isolated kinase domain (**Figure 2—figure supplement 2**). Several factors may contribute to the limited improvement afforded by structure-aware models. Inhibitor binding largely preserves the overall kinase fold, such that structural differences between conditions are relatively subtle. In addition, learned structural representations may emphasize global fold similarity over these local conformational differences, making structure-derived features, including Foldseek tokens (van Kempen et al. 2024), insufficiently discriminative across inhibitor conditions. Finally, the predicted input structures may them-selves attenuate ligand-induced perturbations, further limiting the structural signal available to downstream models. Thus, providing three-dimensional protein structure alone does not necessarily ensure that the ligand-dependent information relevant to mutational fitness is represented or used by the model.

To better understand whether prediction accuracy deteriorates under inhibitor treatment in the same way for different models, we analyzed model behavior from two complementary perspectives: inter-model agreement and sensitivity to ligand context. For the inter-model analysis, Spearman’s correlation coefficients were computed between predictions from different models across all inhibitor conditions (Hou et al. 2026) (**Figure 2**C lower triangle; condition-specific analyses are shown in **Figure 2—figure supplement 3**). Sequence-based protein language models exhibited strong mutual agreement, and a similar pattern was observed among graph-based MPNN-family models, likely reflecting shared training objectives and architectural principles. In contrast, models developed for distinct prediction tasks, including ThermoMPNN (Dieckhaus et al. 2024), PottsMPNN (Birnbaum and Keating 2026) and ProteinEBM (Roney et al. 2025), showed substantially weaker agreement with the remaining methods. We next assessed ligand-context sensitivity by correlating predictions across inhibitor conditions (**Figure 2**C upper triangle). Although the experimentally measured fitness landscapes differed substantially between inhibitors (pairwise Pearson correlations ranging from approximately 0.3 to 0.8; **Figure 2—figure supplement 2**), most models produced highly similar predictions across conditions. In particular, structure-aware protein language models frequently generated predictions with pairwise correlations exceeding 0.9, indicating limited sensitivity to ligand-dependent functional effects. Together, these analyses suggest that current models primarily encode fitness while remaining largely insensitive to ligand-specific functional perturbations. Moreover, the similarities in architectures and approaches render the predictions highly correlated with each other.

### Ligand-aware models do not greatly improve predictions

Next, we evaluated recently developed ligand-aware models, including the protein-chemical language model, ProtLigand, (Kalifa et al. 2025) and graph-based models, LigandMPNN (Justas Dauparas et al. 2025) and LASErMPNN (Fry et al. 2025). ProtLigand connects the structure-aware latents of SaProt (Su et al. 2023) with the hidden embeddings of a ligand from the pretrained ChemBERTa transformer (Chithrananda et al. 2020) using cross attention (Vaswani et al. 2017). This design allows ProtLigand, in principle, to predict mutational effects in the presence of ligand molecules with a log-likelihood ratio trained on the masked token reconstruction task (Meier et al. 2021). In contrast, LigandMPNN extends ProteinMPNN’s (J. Dauparas et al. 2022) architecture through the introduction of k-nearest (k=25) connected residue-ligand graphs which explicitly accounts for the influence of ligand atoms to residue likelihood prediction. Meanwhile, LASErMPNN (Fry et al. 2025)introduces the ligand representation along with protein-ligand interaction graphs into message-passing layers. This was originally developed for zero-shot design of ligand-binding proteins can therefore score a per residue likelihood from a geometric graph of protein-ligand complex. Despite explicitly incorporating ligand information, these approaches also exhibited performance reductions exceeding 50% under inhibitor-treated conditions. Again, this suggests that current ligand representations, or input wild-type structure accuracies, remain insufficient to capture mutation-dependent functional responses (**Figure 2**B). ProtLig remains highly correlated with ligand-unaware models across inhibitor treatments, suggesting only minimal interaction between the specific mutations and the ligand. In contrast, LigandMPNN and LASErMPNN provide more distinct predictions across inhibitor conditions than protein-only models, which indicates potential for these approaches to explicitly model ligand interactions when provided an accurate structural input.

Although overall improvements in prediction accuracy remained modest, ligand-aware models more accurately recovered several experimentally validated resistance-associated hotspots than their corresponding protein-only models (e.g. ProteinMPNN). Under the type I inhibitor crizotinib, LASErMPNN more consistently classified known resistance mutations than ProteinMPNN (**Figure 3**A, upper panel). We attribute that this success is due to the fact that ligand input information is directly encoded and interacts with the associated protein embedding during message passing layers. In particular, mutations at the solvent-front hotspot G1163 (Jean Cui 2013; C. Wang and Lu 2023; Fujino et al. 2019; Recondo et al. 2020); **Figure 3**A), which were predicted as phenotypically neutral or mildly sensitizing by protein-only models such as ProteinMPNN and SaProt, shifted toward experimentally observed resistance phenotypes once explicit ligand information was incorporated. A similar trend was observed for the deeply buried binding-pocket residue L1140, where LASErMPNN consistently predicted mild resistance toward both type I and type II inhibitors, whereas ProteinMPNN remained largely insensitive to ligand context. Structural comparisons (**Figure 3**B) provide a mechanistic explanation for this behavior, local protein conformations surrounding L1140 remain nearly indistinguishable across protein-only predicted structures despite different inhibitor conditions, indicating that protein geometry alone provides insufficient information to distinguish ligand-specific effects. Ligand-aware models were also successful in predicting the profiles for another prominent mutation hotspot (Y1230) (Recondo et al. 2020; Fujino et al. 2021), which directly contacts several type I inhibitors and participates in π-stacking interactions within the binding pocket. However, these improvements were less apparent for type II inhibitors.

**Figure 3.**
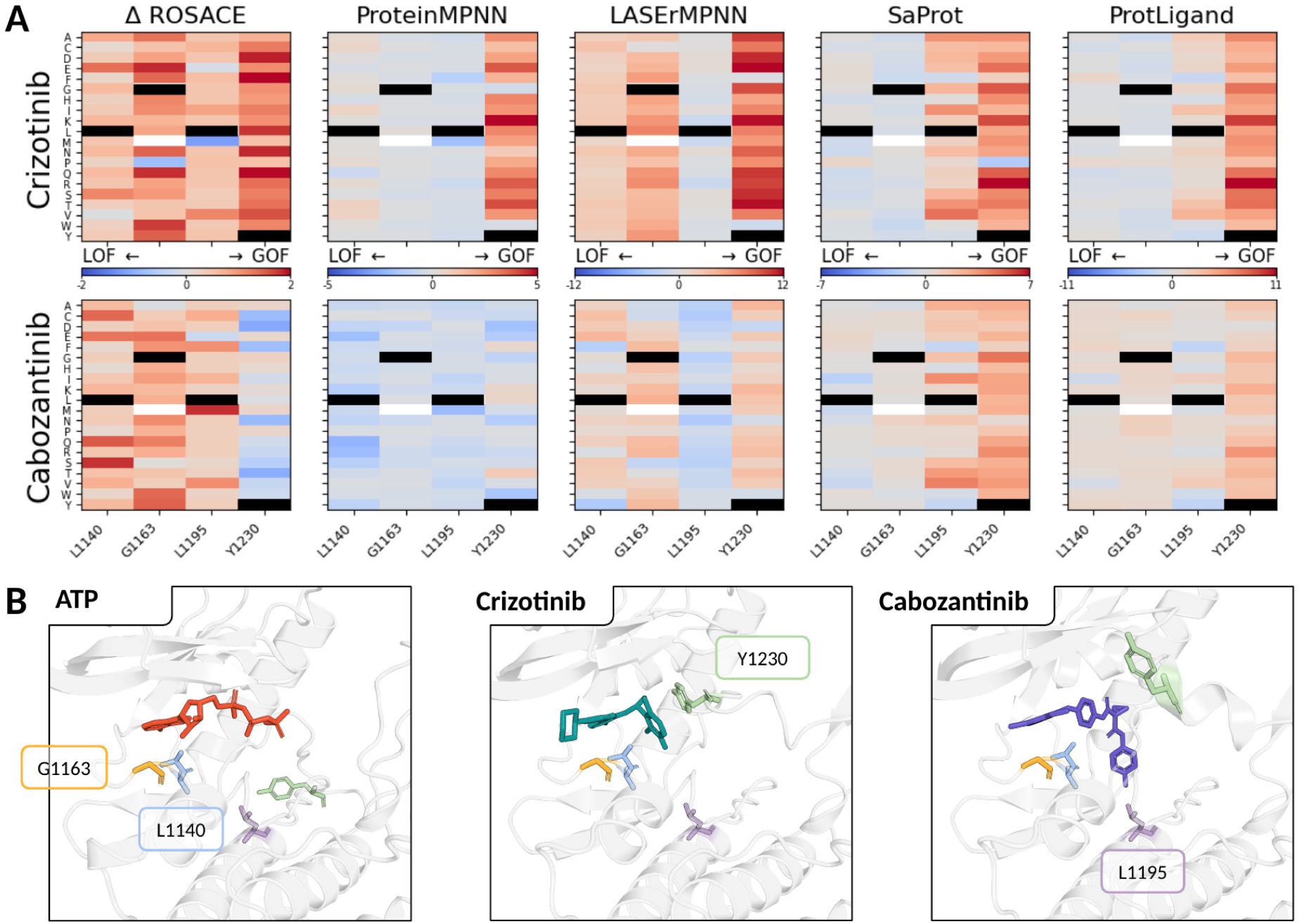
Ligand-aware model predictions at binding-pocket hotspots. A) Models’ predictions on relative mutational effects to the control (predicted with ATP-bound MET kinase) compared against the experimental ROSACE values of mutations (colorbar red to blue indicates gain-to loss-of-functions from control to inhibitor treatments) within the binding pockets where four mutations, L1140, G1163, L1195, and Y1230, are highlighted. Wildtypes are in black and mutations with experimental values missing are colored white. B) AlphaFold3 predicted cofolded structures of MET kinase domain bound to ATP (control, left), Crizotinib (type I, middle) and Cabozantinib (type II, right), which are inputs to the models in A), with color-coded hotspot resistance sites in stick representation.

ProtLigand similarly identified Y1230 as a resistance-associated site under type I inhibitor conditions (**Figure 3**B). However, the performance was similar for SaProt, suggesting that the masked sequence reconstruction objective inherited from conventional protein language models is driving performance. In this case, conditioning protein embeddings on ligand representations through cross-attention does not explicitly optimize protein-ligand interaction representations (Alley et al. 2019; Gordon et al. 2024). Consequently, ligand information appears to exert only a weak influence on the learned embeddings, limiting the ability of current ligand-aware protein language models to capture inhibitor-dependent mutational effects.

In summary, explicit incorporation of ligand information can improve prediction of several clinically relevant resistance hotspots. However, these improvements remained localized to known sites directly adjacent to the ligand (solvent-front, hinge-region, and activation-loop residues). This observation reflects the local nature of the graph construction and indicates that current ligand-aware architectures and/or structure predictions fail to fully capture mutation-dependent protein–ligand interactions. These observations are in agreement with recent work that suggests that many mutational effects arise through long-range allosteric mechanisms rather than large structural rearrangements (Liao and Lehner 2026). It further raises the question of whether current structure prediction models adequately capture the interaction-level perturbations required for downstream protein–ligand prediction. Nonetheless, we continued exploring the utility of predicted mutant structures for estimating ligand affinity.

### Geometric protein-ligand affinity models perform better when single-point mutations are closer to the active site

While in earlier sections we frame the question as reconstructing fitness landscapes under different inhibitor conditions, the task can also be formulated as predicting affinity landscapes of inhibitor molecules to different receptors, in this case different mutants of MET. The conventional approach to this task is geometry-based molecular docking, wherein the binding orientation and affinity between a drug and a protein are estimated based on several physical non-covalent interactions (Kuntz et al. 1982; DesJarlais et al. 1986; Allen et al. 2015). We therefore docked individual inhibitors against mutant structures as our baseline using AutoDock-GPU (Santos-Martins et al. 2021). We also examined deep learning models for drug-target affinity prediction. These deep learning models are typically trained on a broad chemical space and provide a complementary framework for evaluating ligand-dependent mutational effects (**Figure 4**A) (Wei and McCammon 2024; Pan et al. 2026). These approaches represent conceptually distinct approximations of molecular recognition: docking evaluates binding using explicitly defined, simplified physical interaction terms, whereas deep learning affinity models learn statistical representations of protein-ligand interactions from structural and affinity data. Despite this distinction, both approaches remain dependent on the structural representation of the protein-ligand complex. Consequently, when experimentally resolved mutant structures are unavailable, as in most of our benchmark, their sensitivity to mutational effects may ultimately be constrained by the accuracy and mutation responsiveness of the predicted structures used as input.

**Figure 4.**
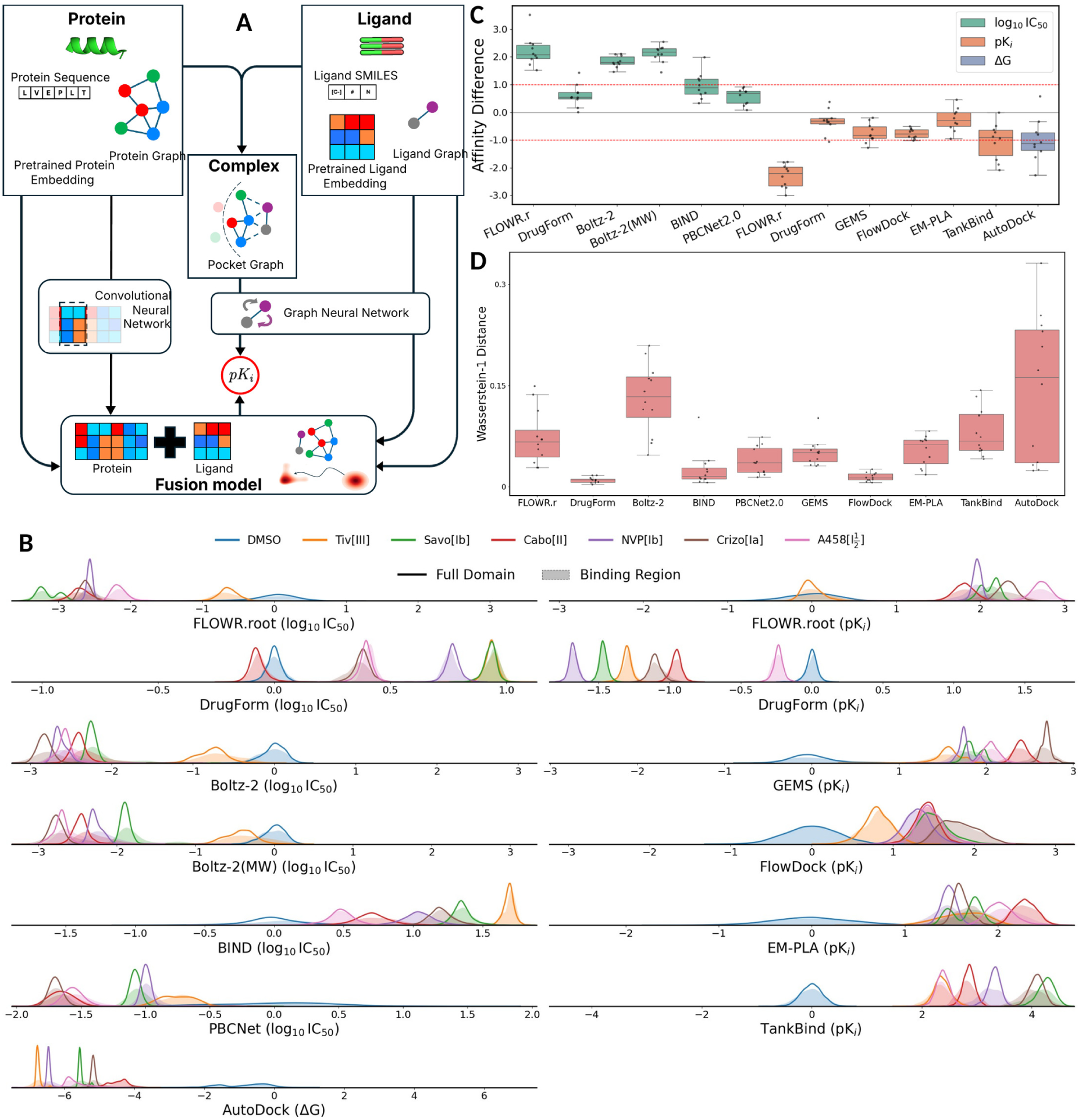
Protein-ligand affinity predictions for MET variants. A) A schematic of current workflow of predicting protein-ligand binding interactions. B) Probability densities of binding affinities predicted by various state-of-the-art deep learning models. Treatments are sorted by their correlations against the control DMSO condition (strongly to weakly correlated from left to right), which is thought to be strongly correlated with the affinity to an inhibitor. The ATP-bound structure was used to represent the experimental control condition. Distributions for all experimental measured variants are plotted in lines and the colored regions correspond to the density of affinities of variants within the binding pocket (which is defined within 5 Angstroms to the ligand). C) Box plots of affinity difference of all inhibitors to Tivantinib (left) and D) Wasserstein distances of inhibitor affinities between variants in the binding pocket and the entire kinase domain (right).

Since many modern affinity models require explicit protein-ligand structures, we used Boltz-2 predicted complexes as inputs to all structure-based methods, which were generated directly with the Boltz-2 structure module. (Passaro et al. 2025; Wan et al. 2026; Bret et al. 2026; Dolorfino et al. 2026) Note that several affinity prediction models (such as TankBind (W. Lu et al. 2022) and FLOWR.root (Cremer et al. 2025)) rely on having a co-folded prediction of ligand and receptor. Instead, they were validated on apo protein structures and ligand SMILES strings as in conventional docking. Our evaluation scheme may therefore introduce underlying inductive biases from Boltz-2 predicted ligand-bound structures during inference. Moreover, recent study by (Feldman et al. 2026) has reported that AlphaFold3-derived structures—and consequently Boltz-2 predictions—are relatively insensitive to single amino acid substitutions due to its dependent on the coevolutionary information derived from multiple sequence alignment augmentation. While most single substitutions do not dramatically reshape proteins, mutational effects in modern structure prediction models are local and their performance deteriorates for strongly perturbative mutations (Zhuoyi Liu et al. 2026). This raises the possibility that mutation-dependent affinity changes may likewise be attenuated in downstream structure-based affinity prediction (further validated with our mutant structures in **Figure 4—figure supplement 1**)

To assess whether these affinity models recover biologically meaningful mutation-dependent effects, we first established two experimentally supported expectations (Estevam et al. 2025). First, the type III inhibitor tivantinib exhibited fitness landscapes that were highly correlated with the DMSO control condition, consistent with comparatively weak target engagement relative to the other MET inhibitors in the panel. (**Figure 1**A, **Figure 2**C and **Figure 2—figure supplement 3**) Accordingly, affinity models should predict substantially weaker binding for tivantinib than for the remaining inhibitors. Second, mutations located within or near the ATP-binding pocket were expected to produce larger perturbations in predicted binding affinity than distal mutations because local structural and interaction changes are more likely to disrupt protein-ligand binding. Consequently, affinity changes associated with binding-pocket mutations should generate broader and more extreme prediction distributions than mutations occurring elsewhere in the kinase domain.

Most structure- and language-based affinity models correctly identified tivantinib as the weakest MET binder, consistent with its DMS fitness landscape closely resembling the DMSO control condition where the control condition was approximated by MET’s binding affinity against ATP (**Figure 4**B). The remaining inhibitors were generally predicted to bind more strongly, with the clearest separation observed for Boltz-2 (Passaro et al. 2025) and FLOWR.root (Cremer et al. 2025) which predicted substantially larger affinity differences between tivantinib and the other tyrosine kinase inhibitors. (**Figure 4**B and C) More importantly, geometry-aware affinity models were considerably more sensitive than transformer-based language models to mutation-induced perturbations within the ATP-binding pocket despite both relying on predicted mutant structures. In particular, Boltz-2, TankBind and FLOWR.root produced broader affinity distributions for binding-pocket mutations than for mutations across the full kinase domain (**Figure 4** C). (Dolorfino et al. 2026). These results indicate that local mutations exert stronger effects on inhibitor binding than distal mutations when evaluated using models capable of representing local conformational adaptations with the potential of modifying the direct interactions with the ligand molecule itself. This result is accomplished in different ways by the two most successful models: Boltz-2 and TankBind incorporate additional single and masked pair representations beyond predicted atomic coordinates, whereas FLOWR.root explicitly refines local protein–ligand conformations during affinity prediction. In contrast, FlowDock (Morehead and Cheng 2024), did not exhibit comparable sensitivity to binding-site mutations, potentially reflecting stronger structural priors inherited from ESMFold-based representations (Lin et al. 2023). Nevertheless, even the best-performing affinity models systematically underestimated the magnitude of mutation-induced affinity changes, likely reflecting either the limited sensitivity of predicted mutant structures, the scarcity of mutation-conditioned supervision in current drug-target affinity training datasets, or the difficulty in scaling between affinity and fitness from the DMS. Together, these findings indicate that current affinity models successfully distinguish broad differences in chemical potency which are likely dominated by properties of the ligand molecules itself, but remain substantially less effective at capturing subtle mutation-dependent perturbations and the corresponding actual protein-ligand interactions (Dolorfino et al. 2026).

To directly compare predicted affinity changes with experimental DMS measurements, we computed the difference in fitness scores between inhibitor-treated and ATP-bound conditions (Δ*f_v_* = *f_v,_*_treatment_ *− f_v,_*_control_), which quantify mutation-dependent changes in inhibitor sensitivity. For this evaluation, we further included two ΔΔG models which were trained or fine-tuned explicitly on experimental measurements of mutation effects (Pires et al. 2015; Yang et al. 2023; Hauser et al. 2018), namely, PremPLI (Sun et al. 2021) and MutDTA (Hu et al. 2025). Across all missense variants, affinity predictions showed little global correlation with experimentally measured fitness changes. (Spearman ρ < 0.2; **Figure 5**A top). Notably, most models perform better with an input of just the binding pocket rather than the full domain(**Figure 5**A; **Figure 5—figure supplement 1**).These observations are consistent with the notion that geometric models predominantly capture local interaction geometry, whereas transformer-based models rely on highly stable latent representations that are only weakly perturbed by single amino acid substitutions.

**Figure 5.**
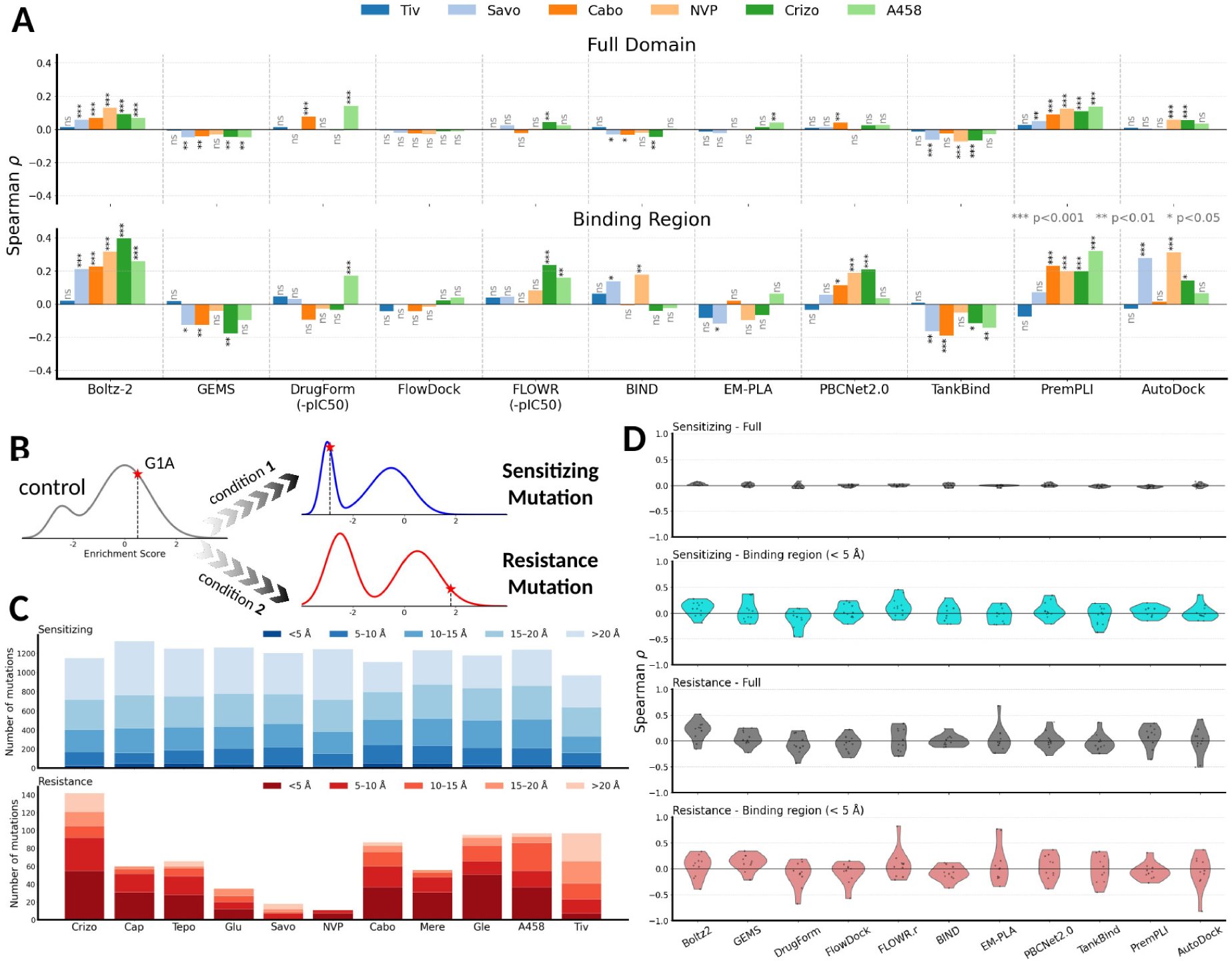
Affinity model performance on resistance and sensitizing mutations. A) Performance of affinity models vs. change in fitness, Δ*f_v_*, for full kinase domain (top panel) and pocket region (bottom panel). Significance levels are indicated as follows: NS (not significant) for *P* >0.05, \**P ≤*0.05, \*\**P ≤*0.01 and \*\*\**P ≤*0.001. B) A schematic to the definition of sensitizing and resistance mutations. Resistance mutations are defined as mutations that exhibit increased enrichment under inhibitor treatment relative to the DMSO control, whereas sensitizing mutations lead to decreased enrichment under inhibitor treatment. C) Spatial distribution of resistance and sensitizing mutations in the dataset. Closer mutated sites were assigned to deeper color. Sensitizing mutations spread over the domain, while resistance mutations are around the cryptic pocket. D) Models’ performance in prediction affinity change in sensitizing (top two panels) and resistance (bottom two panels) subsets averaged over all inhibitor conditions with error bars indicating the standard deviation.

Following recent analysis by (J. Yu et al. 2026), where affinity predictions are evaluated by mutations that preserve experimentally observed binding conformations (i.e., tolerated mutations) and active-site mutations oftentimes impair protein stability (Khersonsky et al. 2018), we classified DMS variants into sensitizing- and resistance-associated mutations. We filtered out non-tolerated mutations in DMSO (**Figure 5**B) and examined the spatial distribution of resistance and sensitizing mutations (**Figure 5**C). A clear distinction can be drawn between the two categories: resistance mutations concentrate at the binding region and sensitizing mutations spread over the full kinase domain. When evaluated separately, all affinity models showed little or no correlation with experimentally measured sensitizing mutations but achieved low to moderate agreement for resistance-associated mutations (**Figure 5**D). Nonetheless, the performance of the models could be significantly improved (from absence of correlation to weakly correlated) when restricted to positions in the binding pocket for sensitizing mutations under certain inhibitor conditions (**Figure 5**D). One possible explanation for this result is that resistance mutations generally preserve the overall protein fold, and, more importantly the key amino acid(s), while perturbing local protein-ligand interactions, making them more compatible with the largely mutation-invariant structures used as model inputs. In contrast, sensitizing mutations may involve larger conformational or stability changes that are insufficiently represented by current structure prediction frameworks. These results suggest that the limited structural sensitivity of predicted mutant complexes constrains downstream affinity models in capturing the full spectrum of mutation-dependent functional effects. Overall, these observations suggest that current *in silico* affinity prediction methods are more effective at identifying mutations that locally perturb protein–ligand interactions within the binding pocket than mutations that induce broader conformational or stability changes across the protein.

Although global agreement between predicted affinity changes and experimental DMS measurements remained limited, multiple affinity models consistently converged on a subset of biologically meaningful resistance-associated hotspots. We therefore identified candidate affinity-sensitive residues by aggregating recurrent outlier predictions across models using a consensus-based ensemble approach (**Figure 6**A). We first examined the contributions of individual models to the recovery of documented resistance hotspots across inhibitor conditions (**Figure 6**B; **Figure 6—figure supplement 1**). For example, Boltz-2 predicted substantial affinity losses for mutations at G1163 and M1211, whereas PremPLI contributed strongly to predictions at D1228. Across models, several clinically established resistance hotspots, including G1163, Y1230, and M1211, were consistently recovered, whereas other documented sites, including L1157, F1134, and G1242, were rarely detected, if at all.

**Figure 6.**
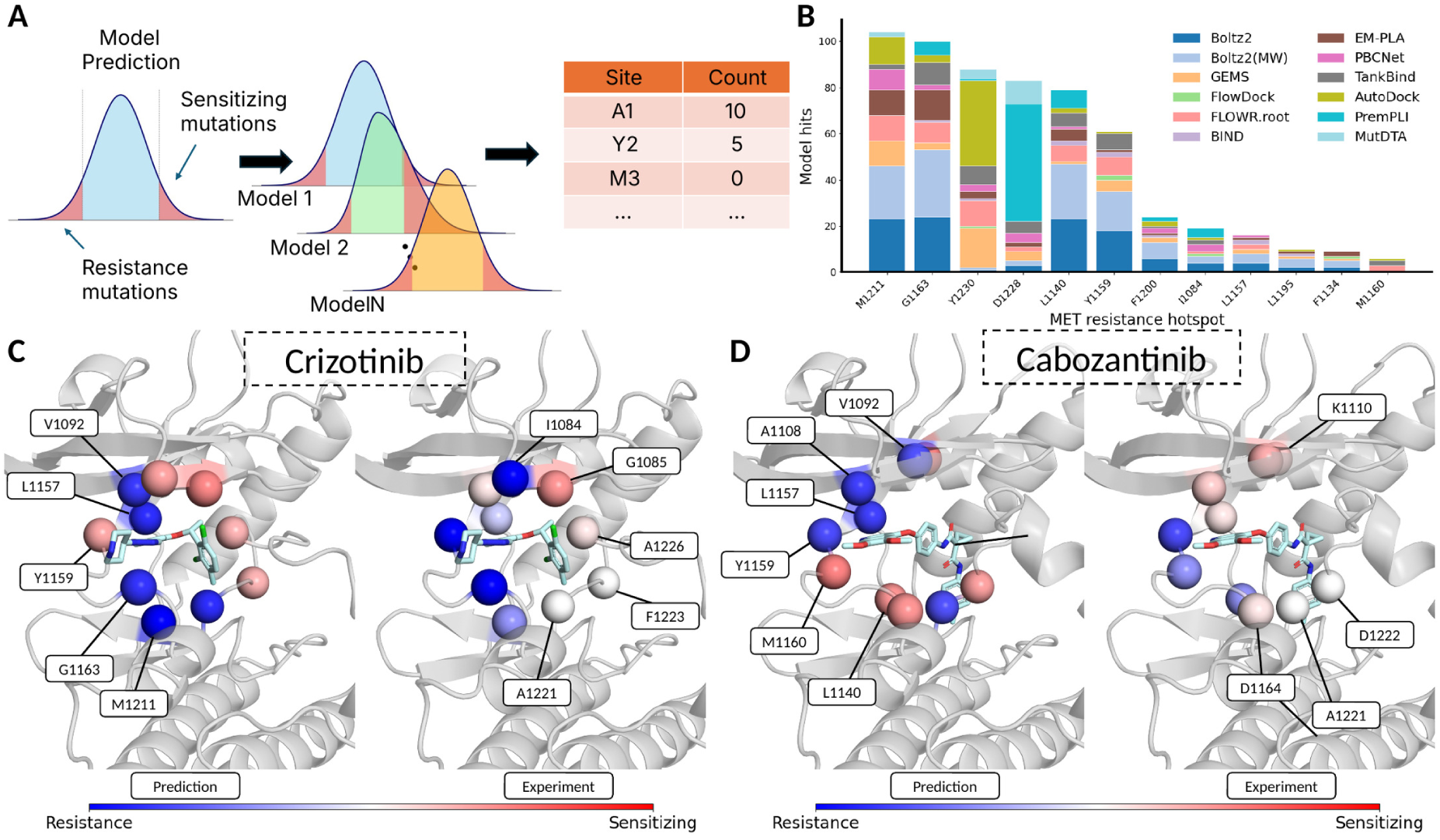
Ensemble prediction of resistance and sensitizing sites. A) A schematic of the ensemble method used for identifying potential resistance and sensitizing positions with affinity predictions made by studied affinity models. B) Models’ ability in recognizing documented hotspot resistance mutations. C-D) Predicted resistance and sensitizing sites to type I inhibitor, crizotinib (subplot C) and type II inhibitor, cabozantinib (subplot D) using ensemble classification of studied affinity prediction models. The results are reported in the left panel and the classifications (to resistance or sensitizing) from experimental measurements of the same positions are in the right panel. Sites are color-coded to blue if they are predicted to be resistance residues and to red for sensitizing residues. Magenta corresponds to critical positions that have both resistance and sensitizing mutations.

We next asked whether cross-model consensus could identify condition-specific resistance and sensitizing sites. We showed the five highest-ranked resistance- and sensitization-associated positions predicted for the type I inhibitor crizotinib and the type II inhibitor cabozantinib (**Figure 6**C). The highest-ranked positions were located within or near the ligand-binding pocket. In particular, mutations at the C-spine residue M1211 and solvent-front residue G1163 were predicted by multiple models, including GEMS, Boltz-2, FLOWR.root, EM-PLA, TankBind, and PremPLI, to substantially reduce crizotinib binding affinity. Likewise, the gatekeeper residue L1157 was repeatedly identified as a resistance-associated hotspot, consistent with its well-established role in regulating access to the hydrophobic back pocket (Tiedt et al. 2011; Hart et al. 2019). For cabozantinib, mutations at the hinge residue Y1159 were predicted to weaken binding and mutations to M1160 within the kinase hinge region, likely reflecting its removal in hydrogen-bonding interactions with bound inhibitors (Damghani et al. 2021)(**Figure 6**D), whereas the same position was less strongly prioritized under crizotinib, ranking ninth among resistance-associated sites (**Figure 6**C). This condition-dependent ranking suggests that the consensus approach can distinguish at least some inhibitor-specific resistance effects.

In contrast, sensitizing predictions showed considerably weaker agreement with experiment. Mutations at the P-loop residue G1085 were consistent with the experimentally measured ROSACE changes and were correctly classified as sensitizing to crizotinib, whereas most other highly ranked sensitizing predictions did not exhibit substantially greater loss of function than in the corresponding DMSO control. Similarly, under cabozantinib treatment, only mutations at the catalytic lysine K1110 among the highlighted sites were consistent with the experimentally observed sensitizing phenotype. Overall, consensus predictions were therefore substantially more robust for resistance-associated mutations than for sensitizing mutations, with the former repeatedly recovering experimentally supported binding-pocket hotspots while most predicted sensitizing sites showed limited agreement with the DMS measurements. Collectively, these analyses suggest that, despite limited global agreement with experimental fitness changes, independent affinity models can recover structurally and clinically relevant resistance hotspots within the MET kinase domain. Cross-model consensus may therefore provide a useful framework for prioritizing candidate resistance mutations for downstream experimental validation.

### Evaluation on diverse kinase-inhibitor DMS datasets yields similar performance

To test whether the lessons from the MET kinase benchmark extended to other deep mutational scanning datasets, we collected other DMS datasets from distinct kinases: SRC (Chakraborty et al. 2024), CDKs (Persky et al. 2020), EGFR (Kim et al. 2024), and FGFRs (Tangermann et al. 2025). While these datasets were measured with different assays, metrics, and data-processing (Table 1), they all fundamentally report on how single amino acid substitutions affect the enrichment of their target when an inhibitor is present.

**Table 1.**
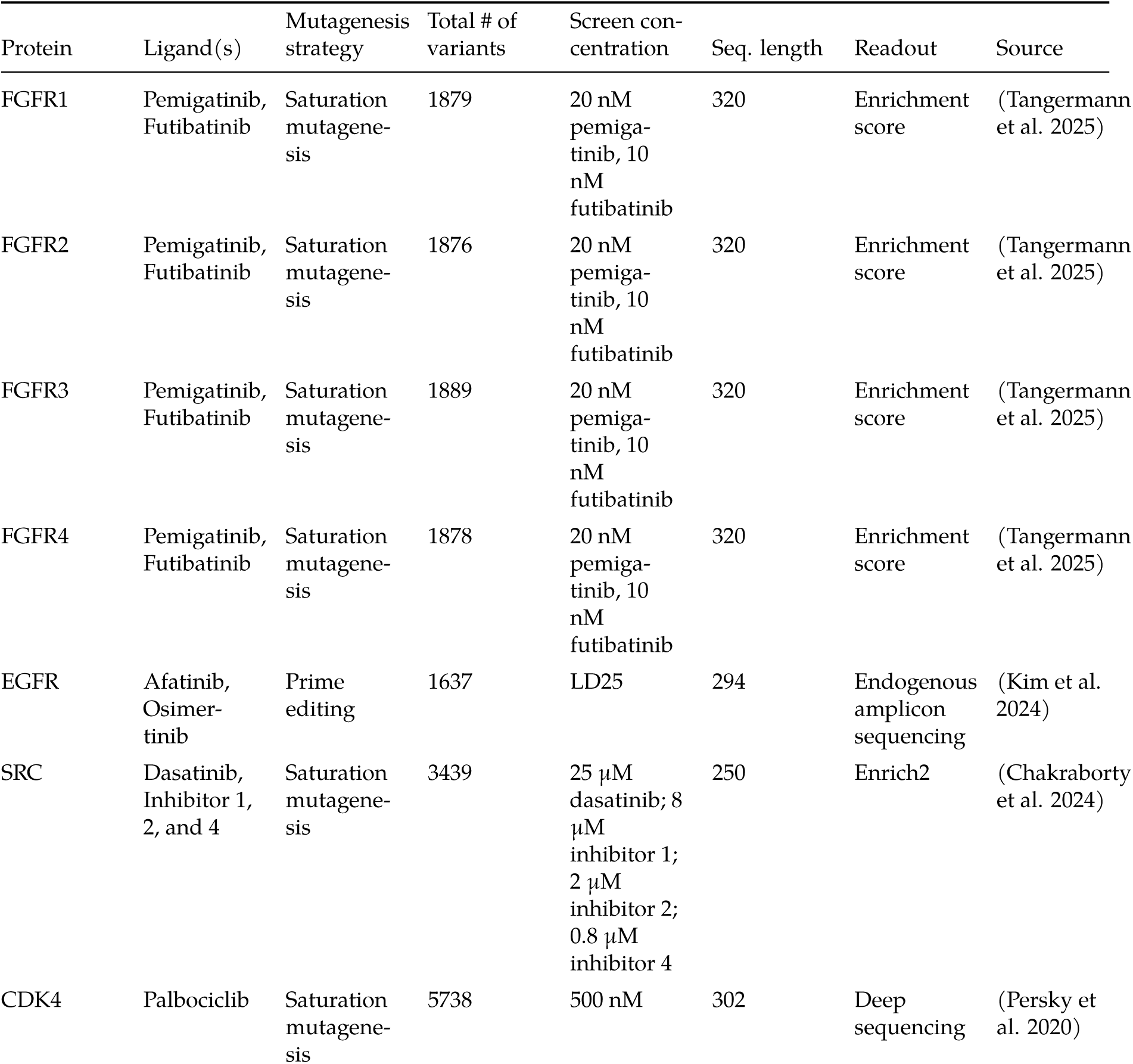
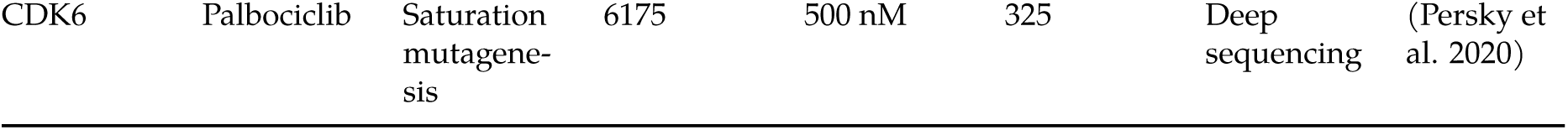
Summary of DMS dataset collected and used for model training.

| Protein | Ligand(s) | Mutagenesis strategy | Total # of variants | Screen con-centration | Seq. length | Readout | Source |
|---|---|---|---|---|---|---|---|
| FGFR1 | Pemigatinib, Futibatinib | Saturation mutagene-sis | 1879 | 20 nM pemiga-tinib, 10 nM futibatinib | 320 | Enrichment score | (Tangermann et al. 2025) |
| FGFR2 | Pemigatinib, Futibatinib | Saturation mutagene-sis | 1876 | 20 nM pemiga-tinib, 10 nM futibatinib | 320 | Enrichment score | (Tangermann et al. 2025) |
| FGFR3 | Pemigatinib, Futibatinib | Saturation mutagene-sis | 1889 | 20 nM pemiga-tinib, 10 nM futibatinib | 320 | Enrichment score | (Tangermann et al. 2025) |
| FGFR4 | Pemigatinib, Futibatinib | Saturation mutagene-sis | 1878 | 20 nM pemiga-tinib, 10 nM futibatinib | 320 | Enrichment score | (Tangermann et al. 2025) |
| EGFR | Afatinib, Osimer-tinib | Prime editing | 1637 | LD25 | 294 | Endogenous amplicon sequencing | (Kim et al. 2024) |
| SRC | Dasatinib, Inhibitor 1, 2, and 4 | Saturation mutagene-sis | 3439 | 25 µM dasatinib; 8 µM inhibitor 1; 2 µM inhibitor 2; 0.8 µM inhibitor 4 | 250 | Enrich2 | (Chakraborty et al. 2024) |
| CDK4 | Palbociclib | Saturation mutagene-sis | 5738 | 500 nM | 302 | Deep sequencing | (Persky et al. 2020) |
| CDK6 | Palbociclib | Saturation mutagene-sis | 6175 | 500 nM | 325 | Deep sequencing | (Persky et al. 2020) |

We followed the same strategy and metrics to assess the ability of different models to predict mutational effects and protein-inhibitor interactions as we used for the MET-inhibitor data. First, we evaluated zero-shot models: ESM2-3B (Lin et al. 2023), SaProt (Su et al. 2023), ProtLi-gand (Kalifa et al. 2025), ThermoMPNN (Dieckhaus et al. 2024), LASErMPNN (Fry et al. 2025) and LigandMPNN (Justas Dauparas et al. 2025). In general, we observed similar performance degradation under non-control conditions as shown (**Figure 7**A). Moreover, among these DMS datasets, language models outperformed geometric models, which is consistent with the lessons from the MET dataset.

**Figure 7.**
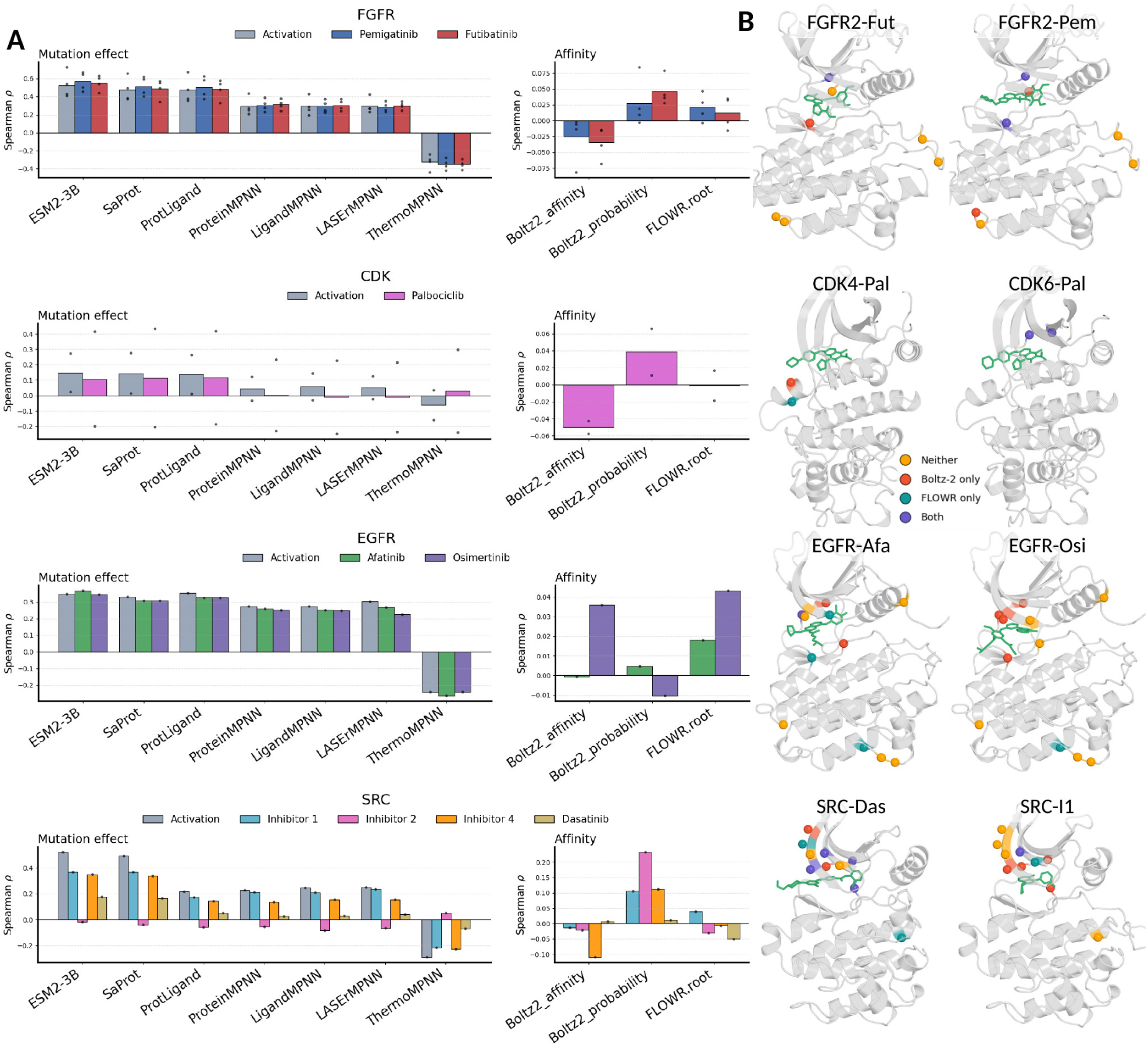
Overview of analyses on kinase-inhibitor deep mutational scanning datasets with mutation effect and affinity prediction models. A) Spearman correlation coefficients of selected deep learning models in predicting mutational effect and affinity against measured enrichment scores. B) Evaluation of Boltz-2 affinity module and FLOWR.root’s performance on documented resistance mutation sites (sphere representations) in colors.

Next, we evaluated affinity predictions from two structure-based models, Boltz-2 affinity module (Passaro et al. 2025) and FLOWR.root (Cremer et al. 2025), which performed best of the models on the MET dataset. Similar to the the MET benchmark, their correlations to enrichment changes over conditions (dtreatment - dcontrol) remained very small (spearman ρ less than 0.1) with Boltz-2 predicted mutant-inhibitor complex structures. We assessed their ability to recover experimentally documented resistance-associated mutations. From over 32 unique resistance sites (60 screenings in total under inhibitor conditions), Boltz-2 flagged 27 of them and FLOWR.root recovered 17 of them (**Figure 7**B), while the two models jointly identified 11 hotspots resistance positions and 25 sites are missed by both of them. In summary, these results demonstrate that the conclusions drawn from the MET benchmark generalize across diverse kinase systems: sequence-based protein language models consistently outperform geometric mutational effect predictors, whereas affinity prediction models remain constrained by mutation-invariant structural representations (Feldman et al. 2026) despite explicitly modeling protein-ligand interactions.

## Discussion

We found that none of the evaluated deep learning models reliably predict mutation-dependent functional responses across inhibitor conditions in MET kinase (Estevam et al. 2025). Although sequence- and MSA-augmented protein language models achieved the strongest overall performance, their predictions remained largely insensitive to ligand context. We attribute the strong performance of protein language models to evolutionary information encoded during large-scale pretraining, which provides informative priors for general mutational effects and likely contributes to their advantage over geometric models that are not directly exposed to comparable sequence variation (Gordon et al. 2024; Hou et al. 2026; Cagiada et al. 2025). However, evolutionary sequence variation and the mutational space accessible through DMS are not equivalent. MSAs reflect substitutions sampled and filtered through evolutionary selection, whereas DMS can systematically interrogate substitutions irrespective of their occurrence in natural sequence space. Inhibitor-conditioned DMS introduces an additional dimension by measuring how the functional consequence of the same mutation changes across chemical environments. Thus, even a rich evolutionary representation of mutational tolerance does not directly encode the mutation-by-ligand interactions required to reconstruct inhibitor-specific fitness landscapes. Consistent with this distinction, pLM predictions remained highly correlated across inhibitor conditions despite experimentally distinct resistance and sensitization landscapes. Models explicitly developed for drug-target affinity prediction showed a complementary limitation, capturing aspects of ligand binding while remaining poorly correlated with mutation-dependent functional responses. Together, these results suggest that current models capture either general mutational constraints or protein-ligand recognition, but not both.

Although structure-aware protein language models can implicitly incorporate spatial information about the inhibitor, they did not consistently improve ligand-conditioned mutational effect prediction over sequence-based models. As an example, in the MET kinase system, type I and type II inhibitors induce only modest global conformational differences, resulting in highly similar structural representations across inhibitor conditions. Consequently, structure-derived embeddings and geometric tokens remain insufficiently discriminative when ligand-specific interactions are absent or only indirectly encoded. While many predicted structures are sufficiently accurate for many downstream structural biology applications at protein-level, the resulting mutation-invariant structural priors in state-of-the-art structure prediction models (Feldman et al. 2026) may obscure the subtle local perturbations that govern inhibitor binding and drug resistance. Consistent with this hypothesis, ligand-aware geometric models, including LigandMPNN, LASErMPNN, Boltz-2, and FLOWR.root, more accurately recovered experimentally validated resistance-associated solvent-front, hinge-region, and gate-keeper mutations than protein-only models. Beyond sequence- and structure-derived representations, recent studies have begun to explore general-purpose large language models, as an additional source of biological and chemical knowledge (Fallahpour et al. 2026; Anand and Goel 2026). Integrating such representations with molecular geometry may provide complementary contextual information that is not readily captured by protein sequence or structure alone, although whether this improves mutation-conditioned protein–ligand prediction remains to be established.

Our results further indicate that incorporating ligand representations through multimodal fusion alone is insufficient to recover mutation-conditioned ligand responses. Although ProtLigand conditions structure-aware protein embeddings on molecular representations through cross-attention, its predictions remained highly correlated across inhibitor conditions and closely resembled those of its backbone structure-aware protein language model, SaProt. One possible explanation is that the masked-sequence reconstruction objective inherited from protein language modeling remains the dominant training signal, while ligand information acts primarily as an auxiliary conditioning signal. In contrast, ligand-aware geometric models explicitly propagate information through protein-ligand contacts. These models showed localized sensitivity only to mutations near the binding pocket. This limitation emerges because the overall training strategy for mutation effect prediction methods emphasize the protein component rather than the ligand contribution. In contrast, affinity prediction models treat these two components equally, but remain biased by ligand features dominating the splitting strategy than protein properties (J. Li et al. 2026). Together, these observations suggest that interaction-aware representations may not emerge simply by combining independently learned protein and ligand embeddings, and motivate training objectives that directly couple sequence variation with ligand-dependent outcomes. (L. Xie et al. 2026; J. Yu et al. 2026) More broadly, mutation effect prediction and mutation-conditioned ligand response prediction should be considered related but distinct learning problems: evolutionary constraints provide strong priors for the former, whereas the latter additionally requires resolving how a specific mutation perturbs molecular interactions within a particular ligand environment.

Importantly, the limitations observed for MET were not specific to a single kinase system. Extending the benchmark to 13 kinase-inhibitor pairs spanning SRC, CDKs, EGFR, and FGFRs reproduced the same broad performance trends across mutational effect and affinity prediction models. At the same time, assembling this expanded benchmark exposed a second limitation: mutation-conditioned kinase datasets remain sparse relative to the large-scale datasets available for related protein fitness and drug–target affinity tasks (Tsuboyama et al. 2023; Notin et al. 2023; Zhihai Liu et al. 2014; T. Liu et al. 2007). Critically, the limitation is not simply dataset size, but also the type and consistency of the available supervision. Protein function is measured using heterogeneous assays, selection schemes, and readouts, making quantitative measurements difficult to compare directly across studies (Anand and Goel 2026). Existing resources such as ProteinGym and FLIP2 partly accommodate this heterogeneity by evaluating models within individual assays or tasks rather than assuming that experimental scores share a common scale (Notin et al. 2023; Didi et al. 2026). While this provides a practical framework for benchmarking, it does not fully address the challenge of learning transferable genotype-phenotype relationships across experiments. This problem becomes more pronounced for mutation-conditioned drug response, where ligand identity and experimental context introduce an additional source of variation. Inhibitor-dependent changes in cellular fitness or enrichment can reflect altered protein-ligand interactions, but can also arise from changes in protein folding, trafficking, catalytic activity, or other determinants of cellular fitness. Without orthogonal measurements that constrain these effects, such as protein surface expression, it remains difficult to determine which molecular consequence of a mutation produces the observed drug-response phenotype. Current drug-target affinity datasets present the complementary problem: they predominantly contain measurements for wild-type proteins and therefore provide limited supervision for learning how sequence variation reshapes protein-ligand interactions.

An ideal mutation-conditioned training resource would therefore align sequence variation, ligand identity and concentration, direct biochemical measurements, and functional phenotypes under standardized experimental conditions. For kinase inhibitors, inhibitor-conditioned DMS experiments could be paired with measurements of protein abundance or surface expression, catalytic activity, direct binding affinity, and clinically validated resistance annotations (Singh et al. 2025; Aldeghi et al. 2019; Saifudeen et al. 2026; Yang et al. 2023). Such orthogonal measurements would help distinguish mutations that directly perturb inhibitor binding from those that alter drug response indirectly through protein stability, expression, or activity, providing the context-aware supervision needed for more reliable prediction of mutation-dependent drug sensitivity and resistance. Meanwhile, synthetic data generated by structure, affinity, or molecular simulation models could help expand sparsely sampled regions of mutation–ligand space. However, such augmentation cannot by itself provide the missing biological supervision: synthetic affinity or structural labels do not necessarily capture downstream effects on protein stability, expression, catalytic activity, or cellular fitness, and may propagate biases already present in the generating models. Synthetic data may therefore be most informative when calibrated against matched experimental measurements rather than used as a replacement for them.

More broadly, our findings indicate that mutation-conditioned drug response represents a distinct challenge from either protein fitness or conventional drug-target affinity prediction. Current protein language models capture strong priors for general mutational effects, whereas ligand-aware and affinity models can identify aspects of protein-ligand recognition and local resistance mechanisms. However, neither capability alone is sufficient to predict how mutational effects change across ligand environments. Addressing this gap will require both models and experimental datasets that explicitly couple sequence variation with ligand-dependent molecular and functional responses. Together, these advances may provide a path toward more reliable prospective prediction of mutation-dependent drug sensitivity and resistance (Urrego et al. 2026).

## Acknowledgements

This work was supported by the NIH (GM145238 to J.S.F.).

## Data Availability

Data analyzed and the corresponding codes in this work is available at https://github.com/fraser-lab/Kinase_Inhibitor_Datasets.

## Methods

### Deep learning methods

In this section we provide a brief description on models evaluated in this work.

#### Structure predictions –

Crystal structures of protein are provided with only protein sequence and ligand SMILES string for AlphaFold3 (https://github.com/google-deepmind/alphafold3) and Chai-1 (https://github.com/chaidiscovery/chai-lab), while steering (via “--use_potentials” flag) is enabled with Boltz-2 (https://github.com/jwohlwend/boltz) predictions. Predictions under template restraints are predicted by specifying “queryIndices” and “templateIndices” under “templates” for AlphaFold3, setting “force” to True and “threshold” to 10 angstrom under “templates” for Boltz-2, and parsing “--use-templates-server” in command line for Chai-1. Since both AlphaFold3 and Boltz-2 allows customized template inputs, we predicted each co-folded MET kinase-inhibitor structure against several templates, and their PDBIDs are 2WGJ, 3DKC, 3DKF, 3DKG, 3QTI, 3RHK, 4EEV, 4R1V, 5T3Q, and 6SDE.

#### Mutational effect predictions –

ESM2 (esm2_t33_650M_UR50D, esm2_t36_3B_UR50D), ISM (ism_t33_650M_uc30pdb) and SaProt (SaProt_650M_AF2) mutational effects are predicted with scripts in SaProt (https://github.com/westlake-repl/SaProt). ESM3 variants (esmc-6B, esm3_sm_open_v1) are predicted with ESM3 codebase (https://github.com/Biohub/esm). Protein language models ProtLigand, PoET and VenusREM are retrieved from https://github.com/kalifadan/ProtLigand, https://github.com/OpenProteinAI/PoET and https://github.com/ai4protein/VenusREM, respectively. In particular, ProtLigand mutational effects are predicted using fine-tuned checkpoint “*final_prot_ligand_model_large.pt”* on mask protein sequence reconstruction of the additional cross-attention head. Both effects from ProteinMPNN and LigandMPNN are predicted with score function in Lig-andMPNN (https://github.com/dauparas/LigandMPNN) with 0.2 Å Gaussian noice checkpoints and autoregressive scoring scheme, respectively. Effects from LASErMPNN (https://github.com/polizzilab/LASErMPNN) are computed by inference logits using checkpoint - “*laser_weights_0p1A_nothing_heldout.pt”*. ThermoMPNN stability predictions are made by https://github.com/Kuhlman-Lab/ThermoMPNN with modification on returning residue probability and corresponding log likelihood. PottsMPNN stability predictions are made by https://github.com/KeatingLab/PottsMPNN.git with default parameters. ProteinEBM (https://github.com/jproney/ProteinEBM) predictions were made with checkpoint *model_6_expert_frozen_1m_md.pt* and timestep 0.1. Models which require geometric information are provided with the co-folded structures predicted by AlphaFold3 and without templates. The complex structure in DMSO condition was predicted with ATP and cofactor Mg^2+^ ion, and the magnesium ion were not provided to all other inhibitor complexes.

#### Drug affinity predictions –

We first predict affinities of all 65,000 missense variants in MET dataset using Boltz-2 with and without molecular weight corrections (via “--use_mw_correction” flag), which also provides a corresponding co-folded structure for each variant-inhibitor pair. These 65k structures are then used as inputs to deep learning models GEMS (https://github.com/camlab-ethz/GEMS), EM-PLA (https://github.com/littlemou22/EM-PLA), Flowr.root (https://github.com/jule-c/flowr_root), PBCNet2.0 (https://github.com/YuJie-0202/PBCNet2.0), FlowDock (https://github.com/BioinfoMachineLearning/FlowDock) after their corresponding data preprocessing workflows. BIND (https://github.com/Chokyotager/BIND), TankBind (https://github.com/luwei0917/TankBind) and DrugForm (https://github.com/drugform/uniqsar), do not require geometric information therefore only variant sequences and ligand SMILES’s are provided. PremPLI mutational effects were predicted using the online web server (https://lilab.jysw.suda.edu.cn/research/PremPLI) All evaluations were made by provided trained model checkpoints, except for MutDTA, which was retrained with details provided in (Hu et al. 2025).

### Docking

The primary tools we used in this work to generate co-folded structures are with a variety of DL models. This becomes cumbersome while inferring ligand-variant interactions, and conversely conventional docking methods (Trott and Olson 2010; Verdonk et al. 2003; Kramer et al. 1999; Allen et al. 2015; Bitencourt-Ferreira et al. 2019) provide a baseline of binding affinity estimations and physically validated binding poses. We then performed docking with GPU accelerated AutoDock4 (Santos-Martins et al. 2021) for computing the binding affinity of all inhibitors toward studied MET variants. These variant models are built with PyMOL mutagenesis wizard using an AF3 (Abramson et al. 2024) predicted apo wildtype structure as template and then preprocessed with Meeko (Santos-Martins et al. 2025). Among the procedures, 662 mutants failed due to severe heavy atom clashes with Meeko and were therefore removed from the analyses.

### Evaluation Metrics

In this work, the performance of a model with predictions *y* to experiment measurements *x* is evaluated primarily with correlation coefficients, including Spearman (Equation 1) and Pearson (Equation 2) correlation coefficients:

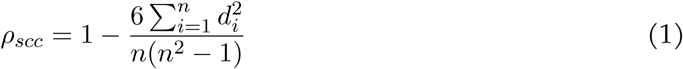

where *d* is the difference between the ranks of *x* and *y*, and *n* is the total number of samples, and

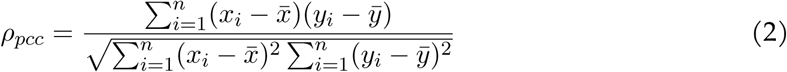

where *x̅* and *y̅* are the mean of samples.

In addition, we compute Wasserstein distance of affinity predictions of variants at full kinase domain and at binding sites. This value can be computed as follow:

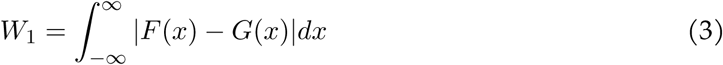

where *F* (*x*), and *G*(*x*) are cumulative distribution functions.

### Mutation Labeling

In this work, we labeled single-point mutations using two methods: 1) binding region mutations and 2) essential mutations. Binding region mutations correspond to mutations that are within 5 Angstroms to any heavy atoms of the ligand, which is a default cutoff used in many deep learning models for defining binding pockets. Essential mutations are variants whose fitness score changes drastically when treated with and without inhibitor molecules. We define mutations that are one standard deviation away from zero in difference scores, *f*_treatment_ *− f*_control_, distribution as essential, in particular, negative difference values refer to sensitized mutation (to the corresponding inhibitor) and positive values indicate resistance mutation (to the corresponding inhibitor).

### Data Collection

Besides the MET dataset, which being the main focus of this work, we collected four additional kinase-inhibitor datasets to increase the generalizability of our model with details summarized in Table 1. These dataset reported either in log-fold change or enrich-2 scores (Rubin et al. 2017).

### Ensemble predictions of low- and high-affinity mutants to inhibitors

To categorize resistance and sensitizing mutations with affinity prediction models, we followed a similar strategy to determine resistance and sensitizing mutations with experimental ROSACE values. In particular, affinity prediction models were trained to predict different experimental observables, such as log_10_ IC_50_ and *pK_i_* = *−* log_10_ *K_i_*. We therefore, negated values predicted in p*K*_i_ which indicates higher values are now predicted to be weaker binder across all predictions made. For each per-inhibitor per-model distribution, the mutations lying at the left tail (i.e., two standard deviations away from the mean) are identified as sensitizing mutations to the corresponding inhibitor and, conversely, mutations are classified as resistance if they are at the right tail.

Next, all models predicted resistance and sensitizing mutations were aggregated. For a position, *p*, its score, Σ is defined in Equation 4.

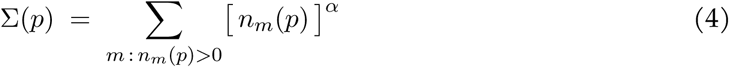

where *m* is affinity model, *n_m_*(*p*) is the number of substitutions at position *p* flagged by model *m*, and the scaling parameter *α* is set to be 0.5. Under this definition, repeated calls from a single model contribute sublinearly, so that concordance across models outweighs repetition within one. Finally, the strong- and weak-binder position scores were ranked independent and the five highest scoring positions were presented and compared against documented and experimentally measured hotspots.

Foundation models capture broad evolutionary and structural patterns,

In addition, some advanced structure prediction frameworks rely on multiple sequence alignment (MSA) preprocessing, which can introduce significant computational overhead and limit scalability. (Perry et al. 2026; Mirdita et al. 2022)

Emerging frameworks further extend these capabilities to machine-learned quantum chemistry, AI-accelerated molecular dynamics (Lewis et al. 2025), and generative modeling of conformational ensembles (Anishchenko et al. 2024; Wankowicz et al. 2024; Lee et al. 2026; Lewis et al. 2025; Jing et al. 2024; Roney et al. 2025).

Sequence-only protein language models serve as an important baseline in this study but do not explicitly represent ligand-dependent conformational states. In contrast, structure-aware deep learning models require complete atomistic protein-ligand complexes as input, motivating us to first assess whether recently developed structure prediction frameworks can provide reliable structural models for downstream benchmarking.

While deep mutational scanning (DMS) experiments provide systematic characterization of mutation-dependent functional landscapes, they are inherently limited to relatively small panels of experimentally tested compounds because each additional inhibitor requires a separate screening experiment.

## Appendix 1

Contents: Appendix 1 text, Equation 4, and Figure supplements (8).

To compare differences in predicted structure to experimental solved structure for MET kinase domain, we reported their global backbone RMSDs against 3DKC in the main text, here we provide more in-depth computations by plotting per-residue backbone RMSDs as of all model predicted ATP-bound complexes (**Figure 1—figure supplement 1**). The same functional regions highlighted in the main text—the P-loop, αC helix, hinge region, and activation loop (A-loop)—are indicated for comparison. Structural deviations were not distributed uniformly across the kinase domain but were strongly concentrated in the A-loop, whereas the P-loop, αC helix, and hinge region generally remained close to the experimentally determined conformation. The reference structure contains two stabilizing substitutions, Y1234F and Y1235D, whereas all predictions were generated using the wild-type MET sequence; therefore, differences within the A-loop should not necessarily be interpreted solely as prediction error. Notably, the models also differed in the secondary structure assigned to this region: AlphaFold3 and Boltz-2 predicted a β-strand-like conformation, whereas Chai-1 produced a more extended/disordered A-loop. Template guidance generally reduced structural deviations, with the largest improvement observed for AlphaFold3, whose global backbone RMSD decreased from 3.68 Å to 1.16 Å and whose A-loop deviation was substantially reduced. In contrast, substantial A-loop differences persisted for Boltz-2 and Chai-1 even with template guidance. Indicating that the predicted complexes reproduce the ATP-binding region and overall kinase fold relatively well, while the conformationally variable A-loop remains a major source of disagreement among structure prediction models.

**Figure 1—figure supplement 1.**
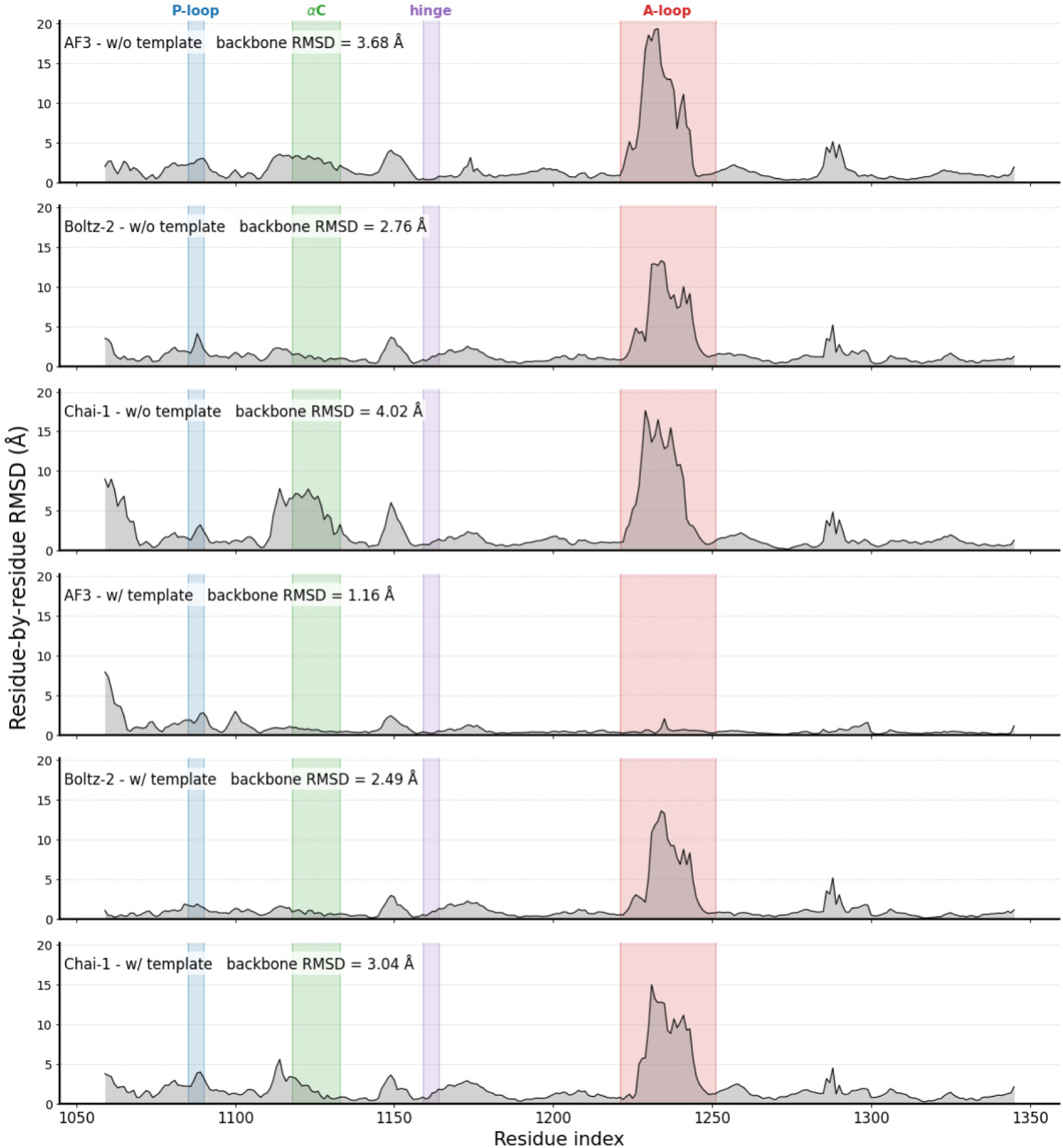
Residue-by-residue backbone RMSDs against reference deposited structure 3DKC. The four regions highlighted in Figure 1 are color-coded here.

As introduced and analyzed in the main text, the mutational effects are predicted by sequence of kinase domains only, which is the region where deep mutational scanning is performed. A complete version of **Figure 2** is presented in **Figure 2—figure supplement 1**. To evaluate the robustness of these models, we further computed the mutational effects with full-length sequences (and the corresponding structures). That means, the input for predicting the functional effect of MET is 1,390 amino acids (vs. 287 amino acids in the main text). Again, complex structures of full Hepatocyte Growth Factor Receptor and inhibitors were predicted with the AF3 model (Abramson et al. 2024). The results are summarized in **Figure 2—figure supplement 2**, where the performances are improved for structure-based and MSA-based models but not for sequence-only ESM models. Evidence can be drawn by computing pseudo log likelihoods (PLL). ((Gordon et al. 2024) Domain sequence yields -1.33 and -1.22 for ESM-650M and ESM-3B respectively, and full sequence has a PLL value of -1.72 and -1.27 for 650M and 3B models. Both values from different ESM sizes suggest the input sequences are of high confidence and therefore the model gains negligible improvement over mutation effect predictions. On the other hand, performances in graph models Protein-, Ligand-, LASEr-, and ThermoMPNN are significantly improved which highlights the necessity of incorporating full descriptions of the object in geometric learning. While full description does seem beneficial for model’s inference, the performances of ProtLigand model (Kalifa et al. 2025) are consistently worsened among all inhibitor conditions (and such phenomena are not observed in its base model, SaProt (Su et al. 2023)). The reason may relate to the PDBbind (R. Wang et al. 2005) dataset used to train ProtLigand’s cross-attention head is composed of truncated structures and, therefore, partial sequences. This observation is in agreement with our analysis and discussion in the Result section on this model as contributions of small molecule inputs in the cross-attention mechanism are not immediately obvious to the model but rather increase the overall trainable parameters in biasing toward true positives to the training datasets.

**Figure 2—figure supplement 1.**
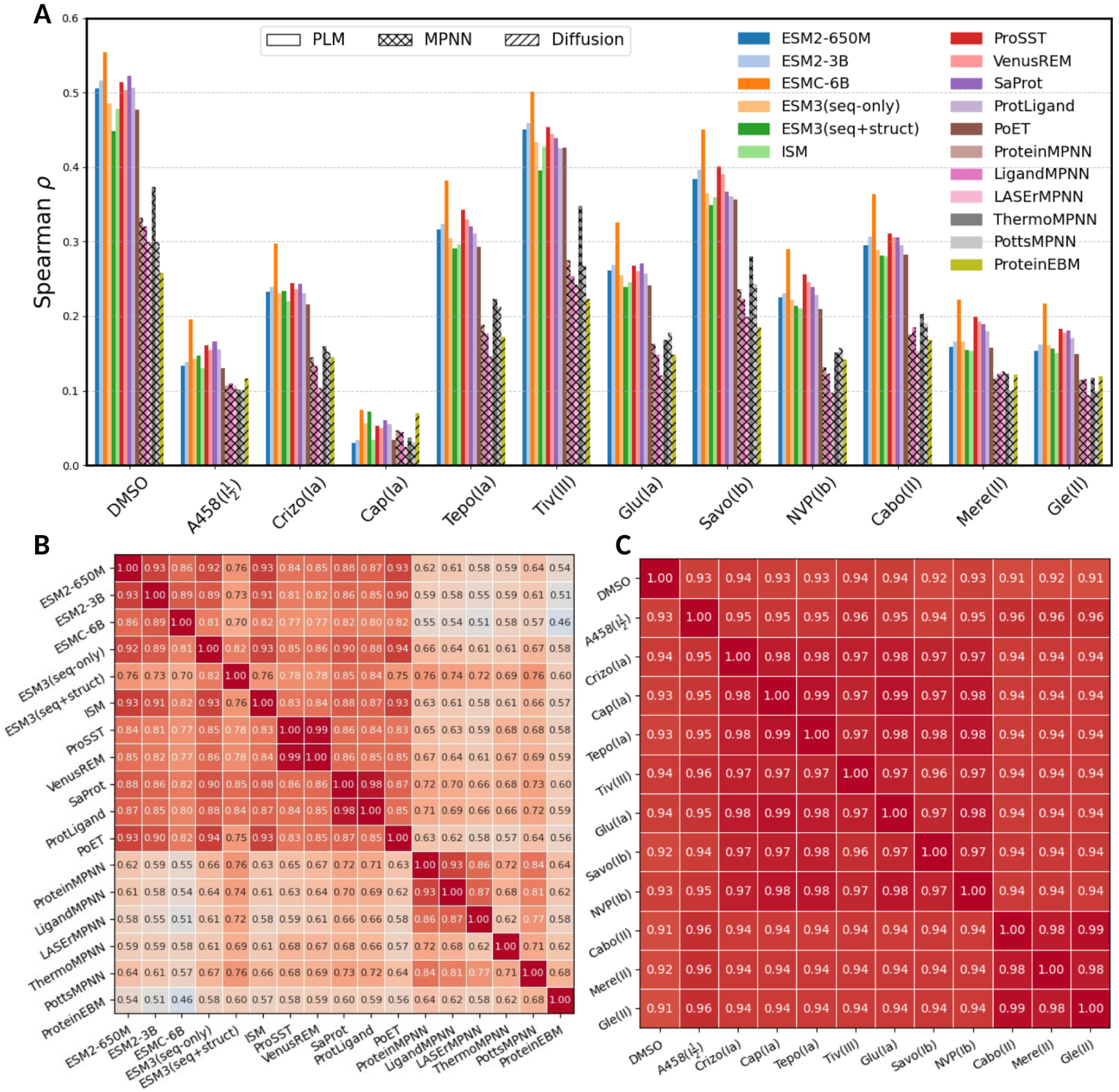
Full benchmark of mutational effect models on the MET DMS dataset. A) Bar plot in Spearman correlation coefficients of all studied models benchmarked on MET DMS dataset. B) Model-to-model and C) treatment-to-treatment Pearson correlation coefficient heatmaps.

**Figure 2—figure supplement 2.**
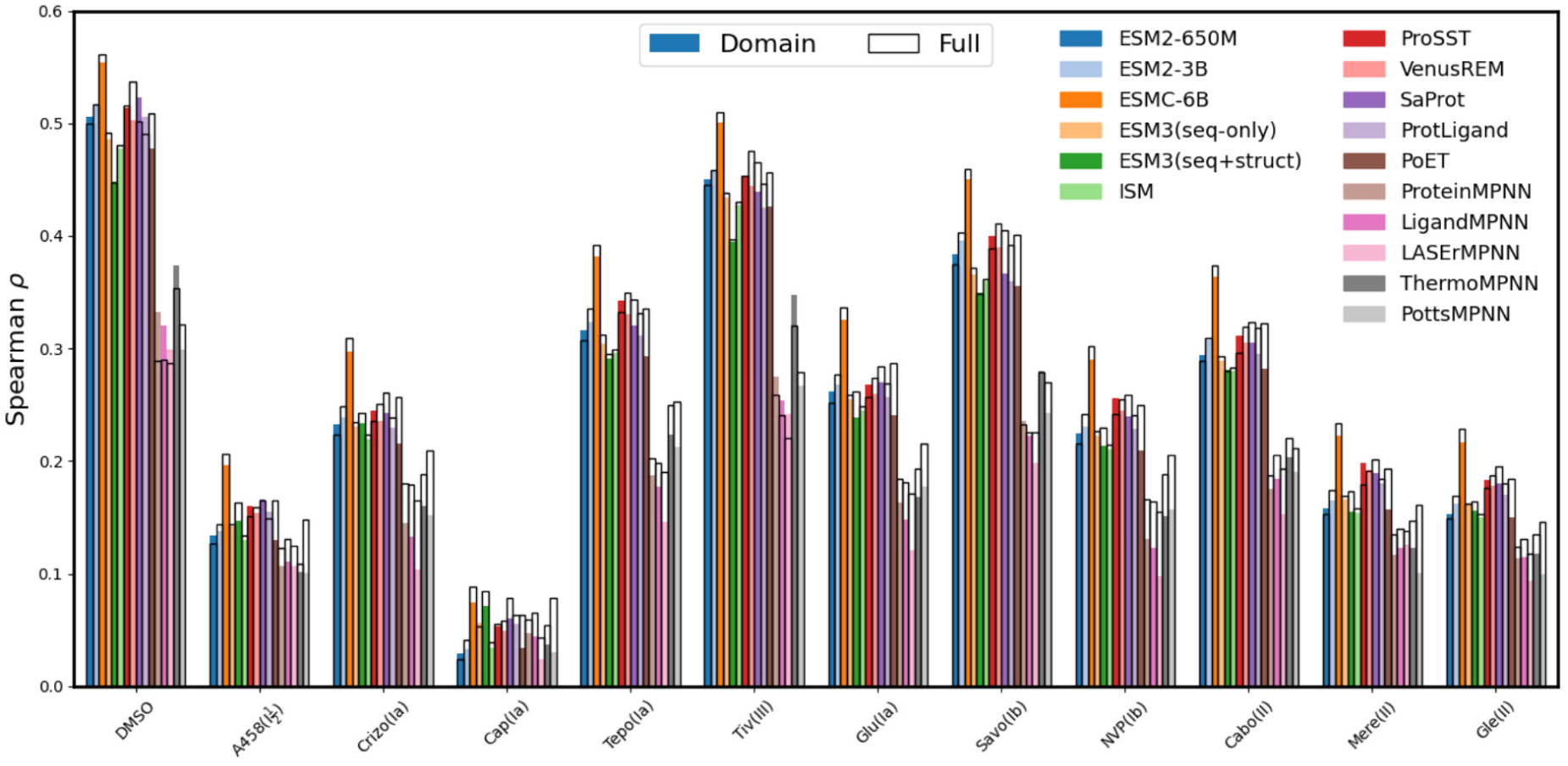
Full-length versus kinase domain sequence inputs. Bar plot of mutational effect predictions with full-length and domain-specific sequence inputs by DL models. Predictions made by domain-specific sequences (and corresponding predicted structure if required) are reported in colored bars and full-length sequences are reported in black frames.

We also report model-to-model and condition-to-condition correlation coefficient heatmaps (**Figure 2—figure supplement 3**) and the averaged maps are reported in main text (**Figure 2**C). As discussed in the main text, MET kinases in different binding modes do not significantly alter model predictions compared to the experimental measures which suggests the information beyond changes in conformation itself is needed for prediction is condition-dependent fitness scores.

**Figure 2—figure supplement 3.**
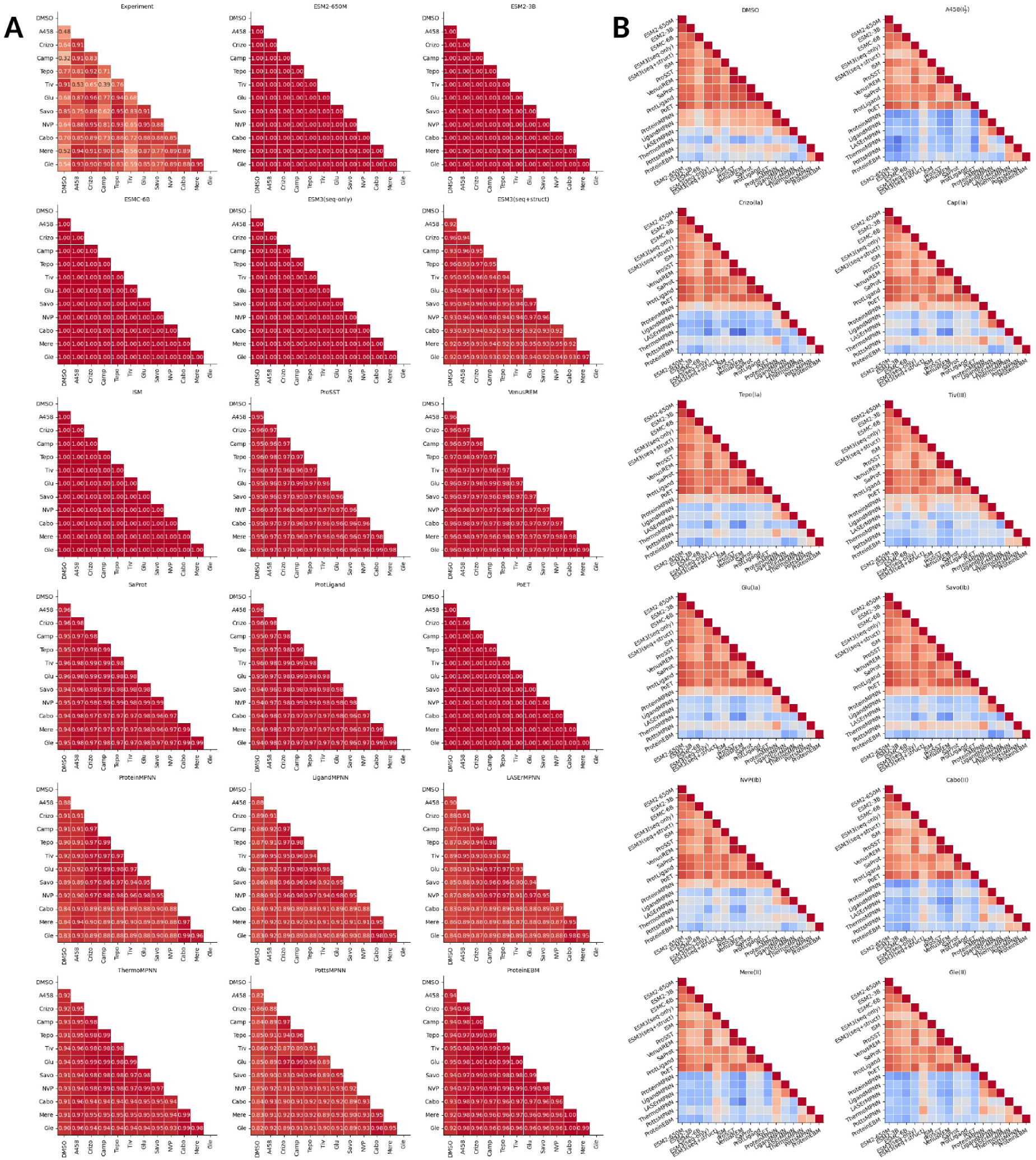
Correlation heatmaps for each model and each condition. Lower triangle heatmaps of Spearman correlation coefficient by deep learning models (A) and by conditions (B).

Several solved structures in the PDB (Berman et al. 2000) were identified and retrieved for template-matching. They are 2WGJ (Jean Cui et al. 2011), 3DKC, 3DKF, 3DKG (Buchanan et al. 2009), 3RHK (Eathiraj et al. 2011), 4EEV (Yan et al. 2012), 4R1V (Dorsch et al. 2015), 5T3Q (Boezio et al. 2017), 6SDE (Collie et al. 2019). These structure templates were provided to AF3, Boltz-2 and Chai-1 for all inhibitor conditions and then used as inputs to two representatives ProSST (M. Li et al. 2024) and LigandMPNN (Justas Dauparas et al. 2025). The relative Spearman correlation coefficients (to predictions made without templates) of full and binding pocket variants are computed and plotted in the top and middle panels of **Figure 1— figure supplement 2**, respectively. The overall effect on having different predicted structures to model predictions is not significant (0.03 change in correlation coefficient in full domain and 0.1 change in binding region). The LigandMPNN model is more sensitive to changes in conformation than the protein language model, ProSST, as the sequence tokens remain invariant in these predictions and they are the dominant component in these protein language models rather than graph neural networks.

**Figure 1—figure supplement 2.**
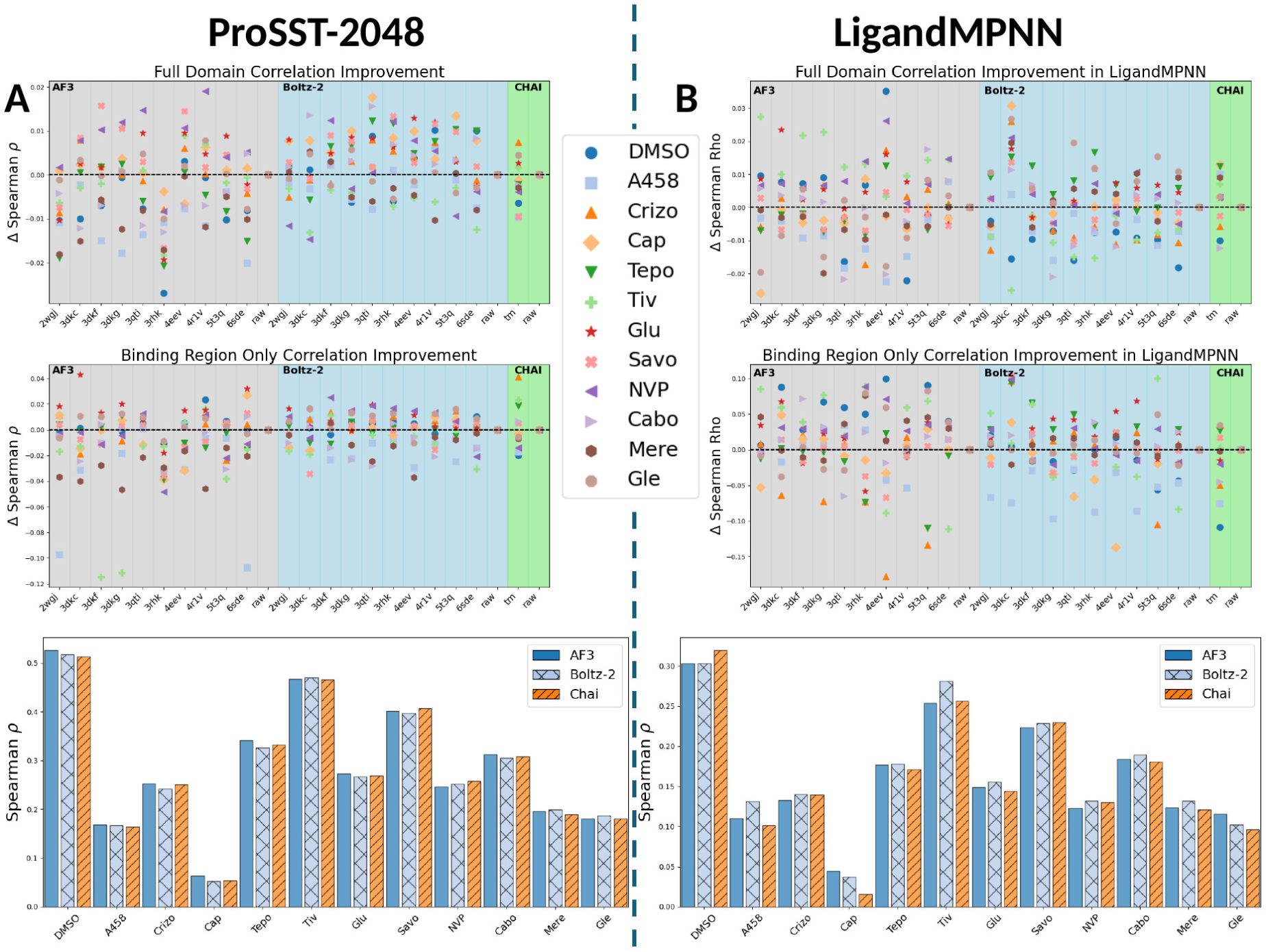
Sensitivity of ProSST and LigandMPNN predictions to the input structure. Investigations on influence of different predicted conformations to protein functional effects in ProSST (A) and LigandMPNN (B). The top panel reports changes in Spearman correlation coefficient with different conformations generated using templates of PDBID in the x-axis (“raw” refers to no template being provided and “tm” means the template search is enabled for structure prediction in Chai-1). The middle panel shows correlation coefficient changes of mutations at the binding region. The lower panel is bar plots directly comparing model predictions with inputs from three models utilized in this work without providing a template.

To evaluate differences in Boltz-2 predicted variant structures, we computed the residue-ligand distances from both Boltz-2 predicted wildtype structures and Boltz-2 predicted variant structures, and these values were plotted against distances from deposited structures. In particular, we highlighted six templates - 2WGJ, 3DKC, 3RHK, 4EEV, 5T3Q, and 6SDE which correspond to inhibitor complexes of MET kinase to Crizotinib, ATP, Tivantinib, Merestinib, AMG-458, and Savolitinib, respectively. For each inhibitor, we predicted one wildtype and all scanned variants in Ref. (Estevam et al. 2025) (5000-6000 structures) with Boltz-2 (Passaro et al. 2025). The results are plotted in **Figure 4—figure supplement 1** suggest Boltz-2 predicted mutated structures show strong correlations to the deposited structures (even than the model predicted wildtype structures) which might be evident for a general limitation of current state-of-the-art models where mutational effects are being largely suppressed.

**Figure 4—figure supplement 1.**
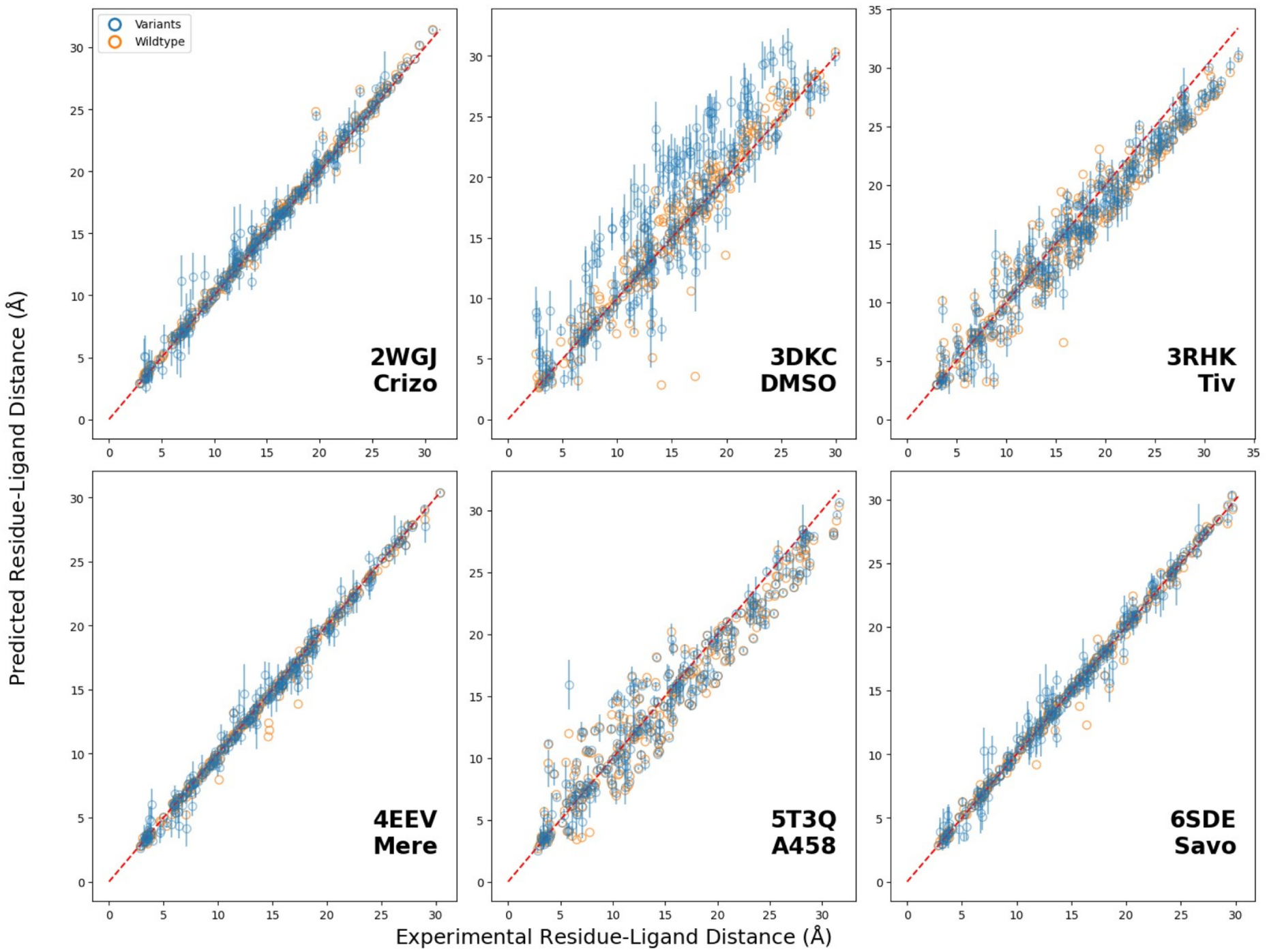
Residue-ligand distances in Boltz-2 predicted wildtype and variant structures. Scatter plot of per-residue residue-ligand distances of deposited structure vs. model predicted structures. The wildtype and mutated structures were predicted by Boltz-2 (Passaro et al. 2025). Orange circles are residue-ligand distances from the predicted wildtype structures at each position and blue circles are the averaged residue-ligand distances from the corresponding predicted variants at that positions with standard deviation in lines.

We predicted all mutagenesis structures with Boltz-2 for its affinity module. We analyzed the variational effect similar to what’s done by (Masters et al. 2025) with complexes whose structure is available in the PDB (Berman et al. 2000). We measured the distances of individual sidechain to ligand’s heavy atom for deposited structure (x axes) and predicted structure (y-axis) (**Figure 4—figure supplement 1**). For predicted structures, we included both wild-type structure (in orange) and mutant structures (in blue; 5000+ for each treatment). Our results suggest mutational effects do not significantly affect the predicted structure while the predictions have preference toward the corresponding wildtype conformation.

With affinity scores predicted from a variety of models, we compared them against the difference of our experimental measured fitness scores between inhibitor treatment and untreated condition (**Figure 5—figure supplement 1**). While most of the models show no significant correlation to experimental measurements, Boltz-2, FLOWR.root, DrugForm, PermPLI, and the conventional method AutoDock show moderate correlations when we narrowed our evaluation scheme to only the binding region, which suggests most of these models emphasize local interactions between the two components, ligand and receptor. Along with stacked histogram in **Figure 6**B over affinity models, we also showcased the same histogram but categorized by inhibitor conditions (**Figure 6—figure supplement 1**), where it shows the resistance and sensitizing position classification is not only model biased, but inhibitor biased.

**Figure 5—figure supplement 1.**
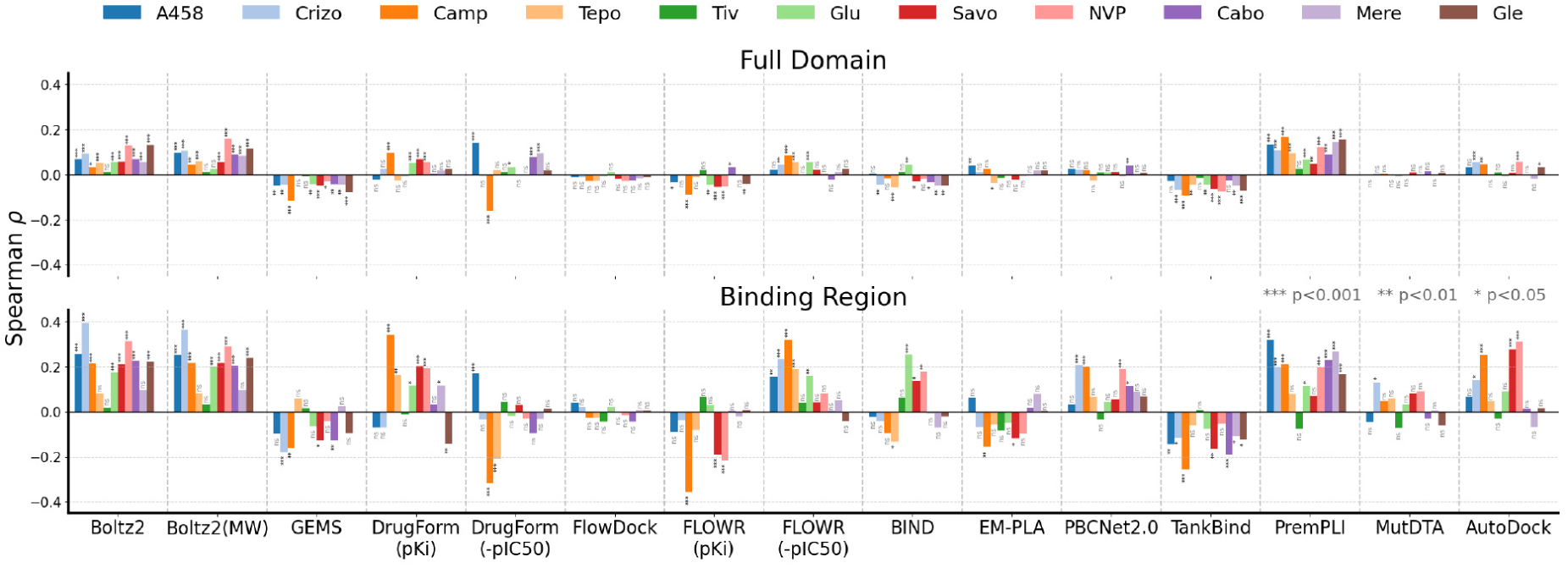
Affinity model performance for all screened inhibitors. Full bar plot of **Figure 5**A which shows the Spearman correlation coefficients of affinity prediction models to experimental fitness scores among full kinase domain (upper panel) and binding region (lower panel) among all screened inhibitors. Significance levels are indicated as follows: NS (not significant) for *P* >0.05, \**P ≤*0.05, \*\**P ≤*0.01 and \*\*\**P ≤*0.001.

**Figure 6—figure supplement 1.**
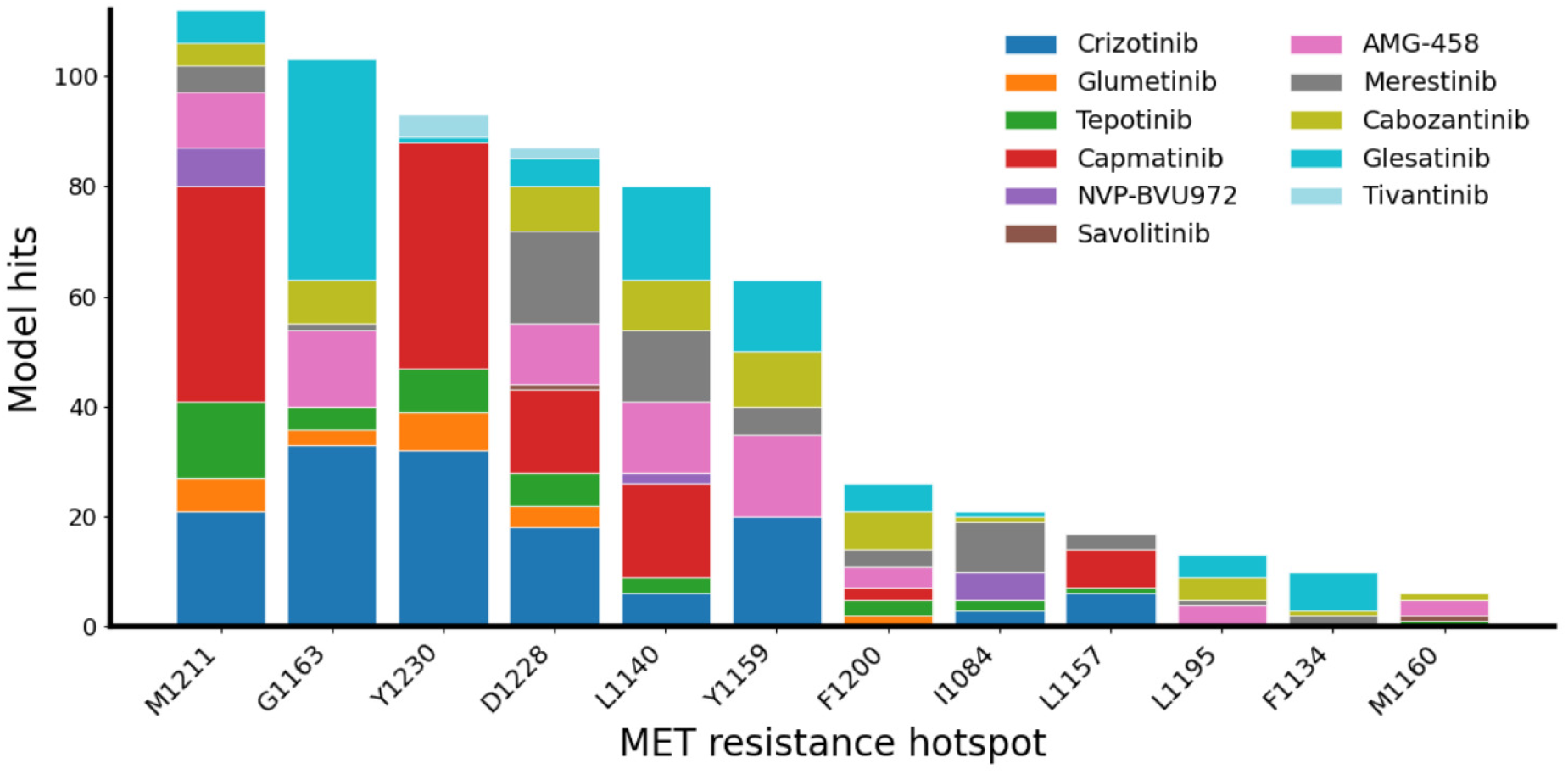
Stacked histograms of inhibitors on documented resistance positions.

